# Neuronal activity-driven O-GlcNAcylation promotes mitochondrial plasticity

**DOI:** 10.1101/2023.01.11.523512

**Authors:** Seungyoon B. Yu, Richard G. Sanchez, Zachary D. Papich, Thomas C. Whisenant, Majid Ghassemian, John N. Koberstein, Melissa L. Stewart, Gulcin Pekkurnaz

## Abstract

Neuronal activity is an energy-intensive process that is largely sustained by instantaneous fuel utilization and ATP synthesis. However, how neurons couple ATP synthesis rate to fuel availability is largely unknown. Here, we demonstrate that the metabolic sensor enzyme O-GlcNAc transferase regulates neuronal activity-driven mitochondrial bioenergetics. We show that neuronal activity upregulates O-GlcNAcylation mainly in mitochondria. Mitochondrial O-GlcNAcylation is promoted by activity-driven fuel consumption, which allows neurons to compensate for high energy expenditure based on fuel availability. To determine the proteins that are responsible for these adjustments, we mapped the mitochondrial O-GlcNAcome of neurons. Finally, we determine that neurons fail to meet activity-driven metabolic demand when O-GlcNAcylation dynamics are prevented. Our findings suggest that O-GlcNAcylation provides a fuel-dependent feedforward control mechanism in neurons to optimize mitochondrial performance based on neuronal activity. This mechanism thereby couples neuronal metabolism to mitochondrial bioenergetics and plays a key role in sustaining energy homeostasis.

## INTRODUCTION

The brain is the most expensive organ to operate, consuming about 20 percent of the body’s resting metabolic energy mostly via the functions of neurons. In intense periods of neuronal activity, this energy demand increases by several fold and imposes an additional burden on neuronal metabolism ^1,2^. Because neurons have limited fuel stores, they meet their instantaneous energy needs by relying mainly on activity-driven ATP synthesis ^3^, fueled primarily by glucose. To ensure a constant supply of glucose when energy demand is high, there is a tight coupling between neuronal activity and glucose metabolism. Neuronal activity transiently increases glucose flux, and leads to rapid glucose utilization for on-demand ATP synthesis via glycolysis and mitochondrial oxidative phosphorylation (OXPHOS) ^3-6^. The biochemical pathways of glucose regulation and flux are generally known; however, the molecular mechanisms regulating ATP synthesis based on glucose availability are poorly understood.

In cells, glucose availability is sensed via the metabolic flux-sensitive post-translational modification O-GlcNAcylation. Nutrient sensor enzyme O-GlcNAc transferase (OGT) catalyzes the reversible addition of a single sugar moiety, O-linked N-acetyl glucosamine (O-GlcNAc), onto serine and threonine residues of proteins. The catalytic activity of OGT is regulated by the intracellular uridine-diphosphate-N-acetylglucosamine (UDP-GlcNAc) concentrations, which fluctuate proportionally in response to nutrient flux through the hexosamine biosynthetic pathway (HBP) ^7-9^. Uniquely, the extent of *O*-GlcNAcylation reflects glucose flux via the HBP pathway. Thus, OGT translates the cellular metabolic state to its >4000 proteins substrates via O-GlcNAcylation ^8,10,11^. Although O-GlcNAcylation has been shown to fluctuate depending on glucose availability in the brain and participate in the regulation of mitochondrial positioning in neuronal axons ^8,12,13^, its role in neuronal metabolism and bioenergetics is unknown.

In this work, we hypothesized that *O*-GlcNAcylation provides glucose flux-dependent feedforward control to optimize mitochondrial performance based on fuel availability, neuronal activity, and energy demand. To test our hypothesis, we used *in vivo* and *in vitro* approaches to demonstrate that neuronal activity upregulates *O*-GlcNAcylation in hippocampal neurons. This upregulation of *O*-GlcNAcylation serves as a signal for mitochondria to sense glucose-flux rate and accelerate ATP synthesis accordingly. To identify the key mitochondrial proteins that are dynamically modified via OGT, we mapped the neuronal mitochondrial *O*-GlcNAcome. We found that *O*-GlcNAcylation regulates mitochondrial plasticity and alters bioenergetics to compensate for the neuronal activity-driven high energy demand. Thus, our study identifies a previously unknown mechanism for the regulation of neuronal energy homeostasis and on-demand ATP synthesis by mitochondrial protein O-glycosylation.

## RESULTS

### Neuronal activity promotes O-GlcNAcylation in neurons

To first determine if neuronal activity directly upregulates O-GlcNAcylation in neurons *in vivo*, we asked how O-GlcNAc levels respond to widespread neuronal activation by inducing seizures in mice via the administration of kainic acid (KA), a kainite-type glutamate receptor agonist ^14^ (Figure S1A). Because KA receptors are particularly abundant in the hippocampus ^15^, we examined sections of the mouse hippocampus for the cellular locus of O-GlcNAc upregulation in the brain by measuring somatic O-GlcNAc levels using immunohistochemistry 1 hour after seizure onset (Figure S1A), having visually verified the induction of the seizure ^16^. We restricted our analysis to the somata of mature neurons by staining for the neuronal marker NeuN ^17^. We identified recently active neurons, which is key for determining rapid adaptations such as transient post-translational modifications, based on expression of the immediate-early gene c-Fos ^18^. Using this approach, we found that seizures enhanced O-GlcNAcylation in neurons throughout the hippocampus, specifically in the dentate gyrus (DG), CA1, and CA3 hippocampal regions (Figure 1A–D, Figure S1B–F). Consistent with elevated neuronal activity, KA administration also enhanced the number of c–Fos-positive hippocampal neurons (Figure 1A–C). To better understand the relationship between neuronal activity and O-GlcNAc levels, we also plotted the intensity of the c-Fos and O-GlcNAc signals for a neuron-by-neuron comparison (Figure 1E). Across the entire population, the O-GlcNAc levels were positively correlated with c-Fos even though O-GlcNAc levels varied independently of c-Fos in some individual neurons. Together, these results indicate that O-GlcNAcylation is upregulated by neuronal activity *in vivo*, and importantly, that this occurs in neurons.

**Figure 1.**
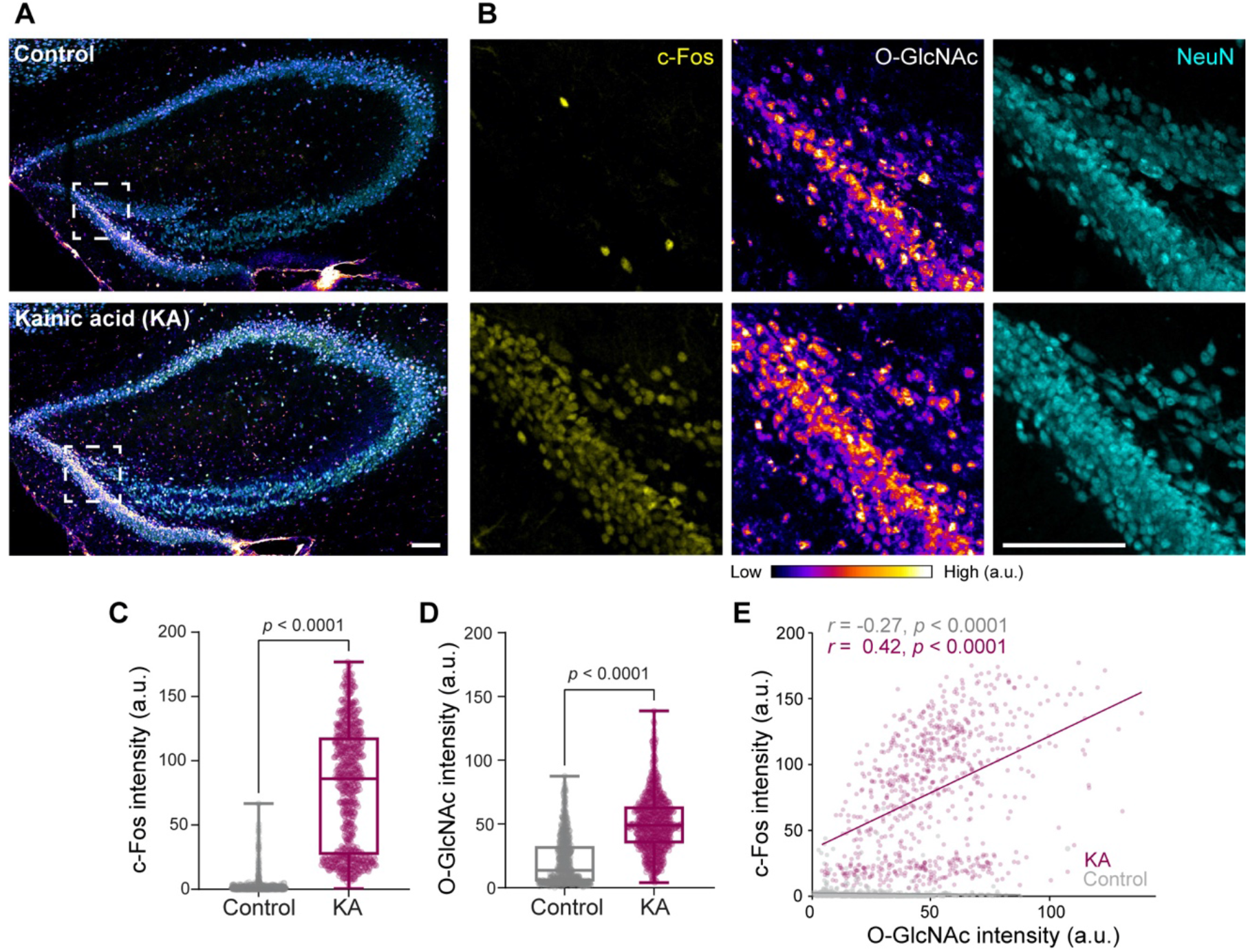
Neuronal activity enhances O-GlcNAcylation in neurons. (A) Representative images of mice hippocampal brain sections after saline (Control) or kainic acid (KA) injections and immunohistochemical staining with antibodies against c-Fos (yellow), O-GlcNAc (RL2, fire LUT), and NeuN (cyan). (B) Magnified images of the highlighted areas (with dashed box) in (A) to show O-GlcNAc and c-Fos fluorescence intensities in the dentate gyrus (DG) hippocampal region in control and KA-treated mice. (C-D) Quantification of c-Fos (C) and O-GlcNAc intensities (D) from NeuN-positive DG neurons from saline (grey) and KA injected (magenta) brains. (E) Scatter plot demonstrating the pairwise correlation between c-Fos and O-GlcNAc fluorescence intensities from individual neurons in DG after saline (grey) or KA (magenta) treatments. The correlation coefficient, *r*, values were quantified by using Spearman nonparametric correlation analysis. For each condition n= 667-720 neurons, 3 mice. Data shown as Min-Max Box-whisker plot with associated p-values (n.s.=p>0.05) (Mann-Whitney U test, Scale bars = 100 µm). See also Figure S1.

### O-GlcNAcylation in neuronal processes is regulated by neuronal excitation

To further characterize the neuronal activity-driven regulation of O-GlcNAcylation at the subcellular level, we carried out immunocytochemistry analysis in disassociated rat hippocampal neurons. First, we examined how O-GlcNAcylation levels change as neurons transition from morphologically mature neurons at 7 days *in vitro* (DIV) to mature network exhibiting spontaneous electrical activity at 14 DIV ^19-21^. Immunostaining neurons for O-GlcNAc at 7 DIV and 14 DIV revealed that neuronal O-GlcNAcylation increased as neurons formed a more mature networks both in the soma as well as in dendrites and axons (Figure S2). Next, we elevated neuronal activity for 6 hours by blocking inhibitory synaptic transmission with the GABA_A_ receptor antagonist picrotoxin (PTX) (Figure 2A–C). Prolonged enhancement of neuronal activity led to a ∼2-fold increase in O-GlcNAcylation within neuronal somas, axons, and dendrites (Figure 2A–G). Importantly, we could reverse this upregulation of activity-driven O-GlcNAc modification using a pharmacological blockade of excitatory synaptic transmission via the selective AMPA receptor antagonist NBQX ^22^ (Figure 2A–G). Thus, our findings demonstrate that increase in excitatory synaptic transmission promotes O-GlcNAcylation in neuronal processes.

**Figure 2.**
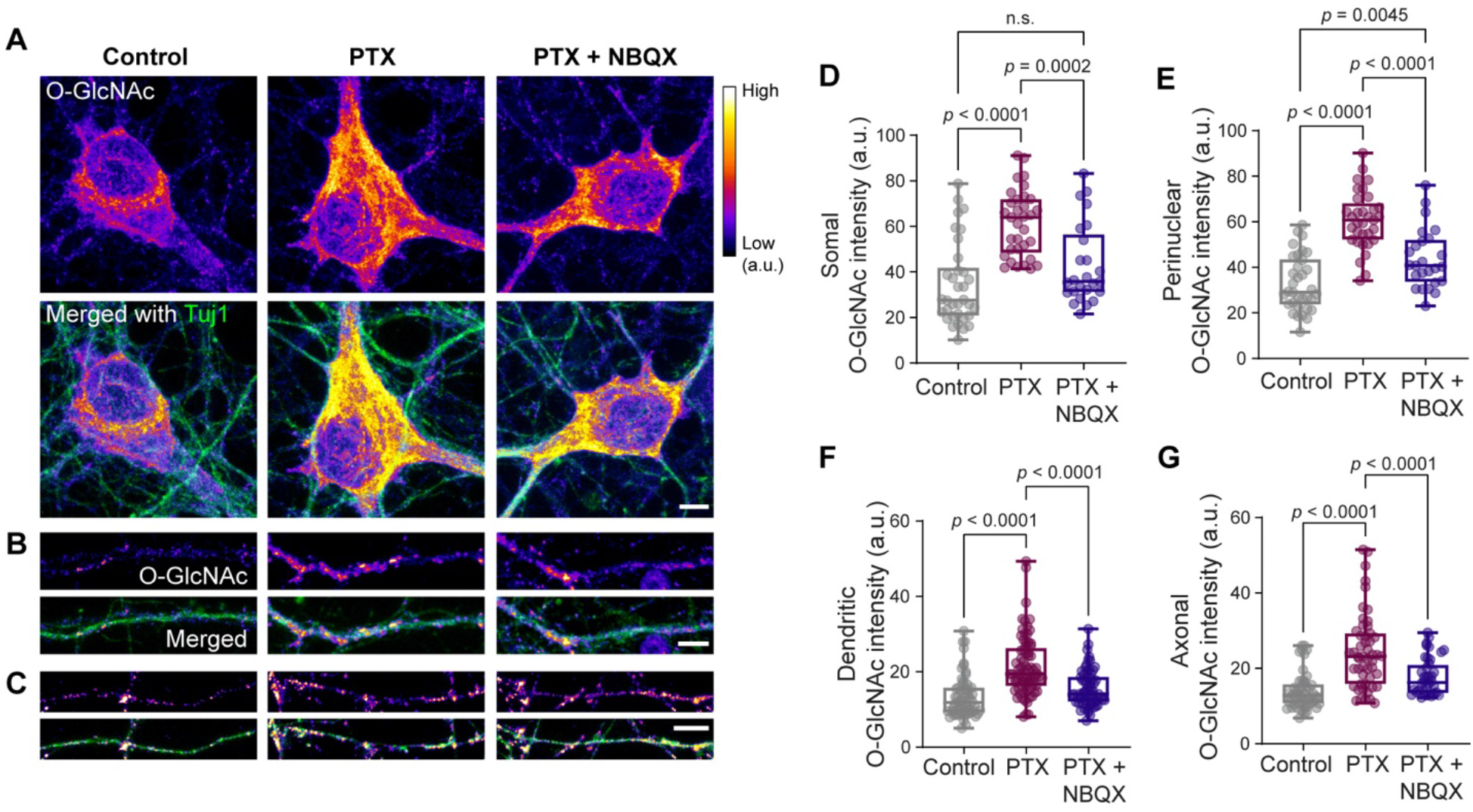
Neuronal excitation promotes O-GlcNAcylation in dendrites and axons. (A-C) Representative images of cortical neuron soma (A), dendrite (B), and axon (C) after 6 hours of DMSO (vehicle control), PTX or PTX+NBQX treatments and immunocytochemical staining with antibodies against Anti-III Tubulin (Tuj1) (green) and O-GlcNAc (RL2, fire LUT). (D-G) Quantification of O-GlcNAc intensities from somal (D), perinuclear (E), dendritic (F), and axonal (G) regions from cortical neurons after indicated treatments. n = 25-34 neuronal soma, 58-73 dendrites, 36-50 axons from three independent experiments. Data are shown as a Min-Max Box-whisker plot with associated p-values (n.s.= p>0.05) (one-way ANOVA with post hoc Tukey’s multiple comparison test, Scale bars = 5 µm). See also Figure S2.

### Neuronal activity-driven O-GlcNAcylation targets mitochondria and regulates mitochondrial bioenergetics

We next set out to identify the intracellular localization of O-GlcNAcylated proteins in neurons, focusing initially on mitochondria. We hypothesized that the metabolic sensor O-GlcNAcylation regulates mitochondria to support elevated neuronal energy demands. Therefore, we examined the subcellular localization of O-GlcNAc in neuronal processes together with the pan-mitochondrial marker protein Tomm20 (Figure 3A–B). When neuronal activity was elevated for 6 hours by PTX treatment, the co-localization of O-GlcNAc with mitochondria also increased in dendritic and axonal processes. This co-localization was reversed when excitatory synaptic transmission was blocked by NBQX treatment (Figure 3A–D). To further evaluate mitochondrial O-GlcNAc levels, we isolated crude mitochondrial and cytosolic fractions from cultured neurons and analyzed them by immunoblotting. In these samples, PTX treatment predominantly increased O-GlcNAcylation of mitochondrial proteins compared to cytosolic proteins (Figure 3E–G). The increase in mitochondrial O-GlcNAcylation was not due to alterations of endogenous OGT or OGA protein levels (Figure S3A). The transient inhibition of OGT activity reduced O-GlcNAcylation in neurons (Figure S3B, 4A), which further supports the dynamic nature of O-GlcNAc modification.

**Figure 3.**
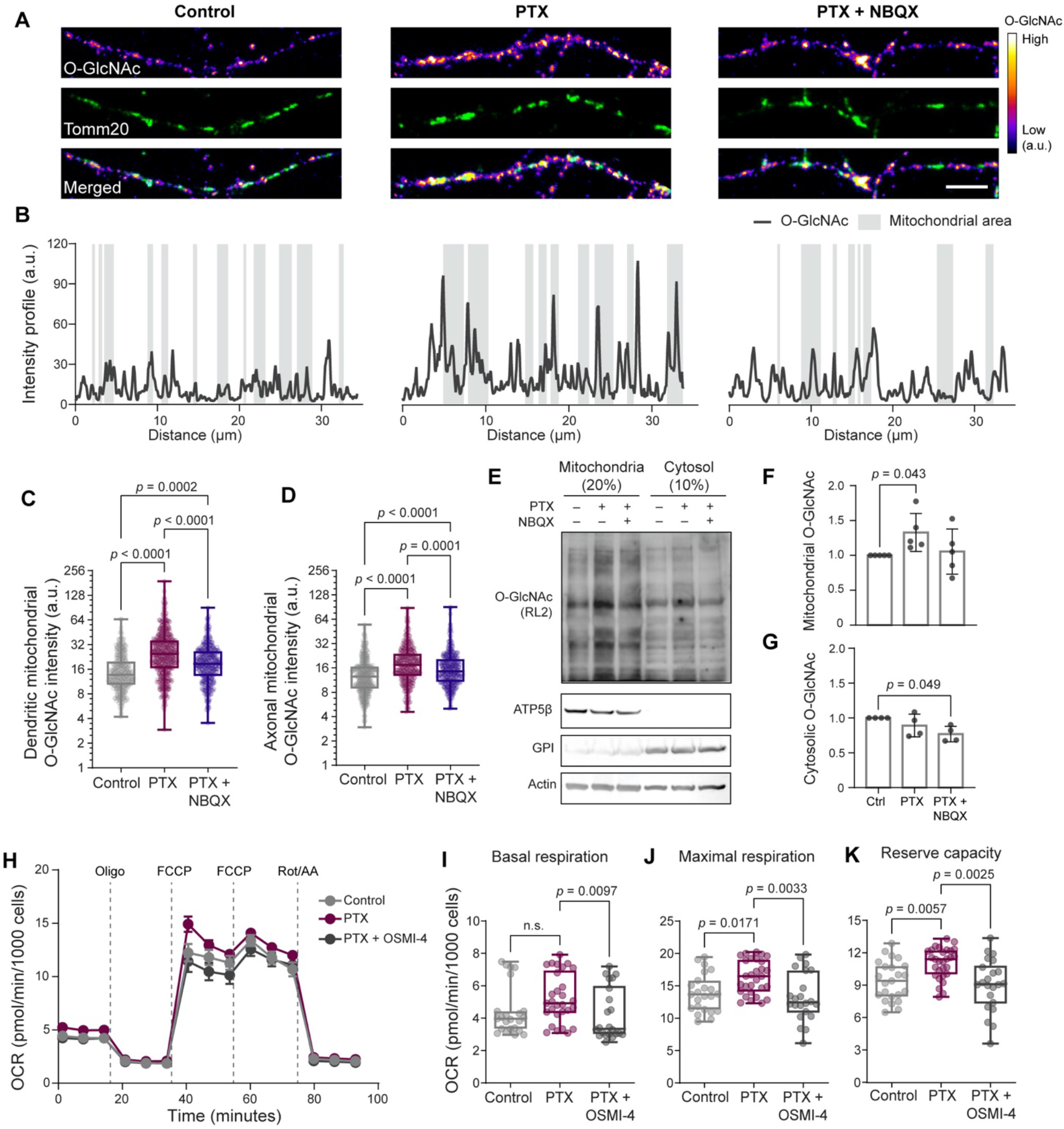
Neuronal activity-driven O-GlcNAcylation regulates mitochondria. (A) Representative images of neuronal processes after 6 hours of DMSO (vehicle control), PTX or PTX+NBQX treatments and immunocytochemical staining with antibodies against Anti-Tomm20 (green) to label mitochondria and anti-O-GlcNAc (RL2, fire LUT). (B) Immunofluorescence intensity of O-GlcNAc and mitochondria measured along the length of neuronal processes shown in (A). (C-D) The total intensity of O-GlcNAc immunofluorescence at the dendritic and axonal mitochondrial compartments after indicated treatments. n = 1000-1281 mitochondria analyzed for each group from three independent experiments. Data are shown as a Min-Max Box-whisker plot with associated p-values (one-way ANOVA with post hoc Tukey’s multiple comparison test, Scale bars = 5 µm). (E-G) Mitochondrial and cytoplasmic protein O-GlcNAcylation level changes. Mitochondrial and cytosolic fractions, obtained from cortical neurons treated with DMSO (vehicle control), PTX, or PTX+NBQX treatments for 6 hours, were separated by SDS gel electrophoresis and probed with anti-O-GlcNAc antibodies (RL2), anti-ATP5β (mitochondrial marker), anti-GPI and anti-actin (cytosolic markers) antibodies. The total intensity of the O-GlcNAc immunoreactive bands was normalized to the intensity of ATP5β (F) or actin (G) bands for each lane. O-GlcNAc levels in control cells were set as 1, and fold changes in response to indicated treatments were calculated. n = 4-5 independent experiments. Data are shown as mean values ± SEM with associated p-values (one-way ANOVA with post hoc Kruskal-Wallis multiple comparison test). (H) Mitochondrial oxygen consumption rate (OCR) measured from cultured cortical neuron cultures using the “mito stress test” protocol after treatments with DMSO (vehicle control), PTX, or PTX+OSMI-4 for 6 hours. (I-K) Basal (OCR_basal_) and maximal respiration (OCR_FCCP_) and reserve capacity (OCR_FCCP_ - OCR_basal_) for indicated treatments. Mean ± SEM for each time point, n= 24-32 wells per conditions from three independent experiments. Data are shown as a Min-Max Box-whisker plot with associated p-values (n.s.= p>0.05) (one-way ANOVA with post hoc Tukey’s multiple comparison test). See also Figure S3.

Because neuronal activity regulates mitochondrial O-GlcNAcylation, we next assessed the impact of O-GlcNAcylation on neuronal bioenergetics. To measure mitochondrial respiratory activity and the glycolytic rate, we cultured primary cortical neurons under physiological glucose levels and performed a respirometry analysis at 12–15 DIV, when the neuronal networks exhibit spontaneous electrical activity (Figure S3C). After each respirometry measurement, we quantified neuronal enrichment in primary neuronal cultures, and differentiated neurons from the glial cells, by co-staining each well with the mature neuron marker NeuN and the nuclear dye NucBlue (Figure S3D). This staining revealed that neurons represent 60–80% of the cellular population in our cultures (Figure S3D–E). We did not observe a change in mitochondrial respiration that would drive basal mitochondrial ATP synthesis. Next, to examine maximal respiratory capacity of mitochondria, we stimulated uncoupled respiration using carbonyl cyanide-4-(trifluoromethoxy)phenylhydrazone (FCCP), and found that neuronal activity increases mitochondrial respiratory capacity (Figure 3H–K). Elevating the neuronal activity with PTX in the presence of the OGT inhibitor OSMI-4 (Figure S3C) reduced both the basal and maximal mitochondrial respiratory capacity (Figure 3H–K). OSMI-4-treated neurons also displayed a reduced glycolytic capacity (Figure S3F–H). These results confirm the involvement of OGT and mitochondrial O-GlcNAcylation in neural activity-driven metabolic flexibility.

### Acute neuronal stimulation triggers glycolysis and mitochondrial O-GlcNAcylation

We next sought to establish the extent to which neuronal activity-dependent glucose metabolism might contribute to mitochondrial O-GlcNAcylation. To monitor the metabolic changes that occur following neuronal activity, we measured fructose 1,6-bisphosphate (FBP) levels in neuronal processes using HYlight, a fluorescent sensor that reports glycolytic dynamics ^23^ (Figure 4A). Because neuronal activity causes transient dips in intracellular glucose levels and also transiently increases neuronal glycolysis ^4^, we predicted that FBP levels would decrease as neuronal activity increases. Instead, measurements from neuronal processes revealed that even persistent action potential (AP) firing (10Hz, 600 AP) produced only small changes in cytosolic FBP levels. As a control, we confirmed that our stimuli induce neuronal activity using the Ca^2+^ sensor GCaMP6s (Figure S4A–B). Therefore, either neuronal activity does not significantly alter glycolysis or increases in demand are accompanied by activity-driven glycolytic flux. When the production of FBP was blocked by exchanging glucose for 2-deoxyglucose (2DG), the FBP levels slowly declined (Τ = 3.007 min, Figure 4B–D, S4C). Neuronal stimulation under these conditions caused a more rapid decay in FBP levels (Τ = 1.007 min), which implies that neuronal activity causes a further increase in FBP consumption.

**Figure 4.**
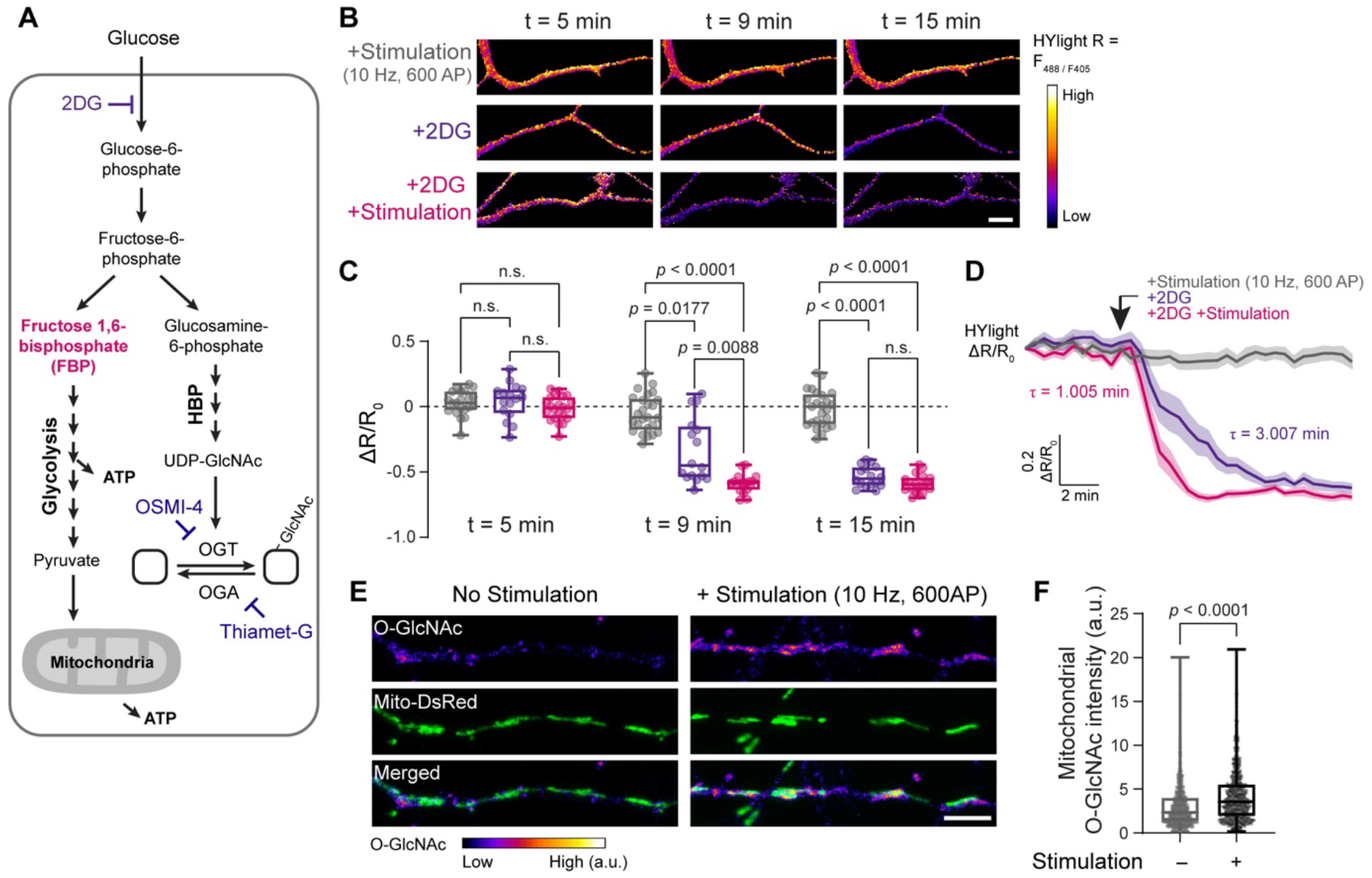
Glycolysis and O-GlcNAcylation rise upon neuronal stimulation. (A) Schematic representation of cellular glucose utilization pathways: glycolysis and hexosamine biosynthetic pathway (HBP). Genetically encoded sensor HYlight measures fructose1,6-bisphosphate (FBP) (magenta), the product of the committed step of glycolysis. Rate limiting steps and the inhibitors used in this study are also indicated: glycolysis inhibitor 2-deoxyglucose (2DG), O-GlcNAc transferase (OGT) inhibitor OSMI-4, O-GlcNAcase, (OGA) inhibitor Thiamet-G. (B) Representative ratiometric HYlight (fire LUT) images from neuronal processes, before (t= 5 min), and after (t= 9 and 15 min) field stimulation (10Hz, 600AP) either with or without 2DG treatments. (C-D) The normalized HYlight emission ratio (R) induced by 488nm and 405nm excitations (ι1R/R_0_) at indicated time points, obtained from cortical neuron traces demonstrated as in (D) (gray: control with field stimulation; purple: only 2DG treatment; magenta: 2DG treatment with field stimulation). The arrowhead indicates the initiation time point of 2DG treatment and the field stimulation (10Hz, 600AP). Data are shown as mean values ± SEM (D) and a Min-Max Box-whisker plot (C) with associated p-values (one-way ANOVA with post hoc Tukey’s multiple comparison test), n = 7-11 neurons from three independent experiments. Scale bar = 10 µm. (E-F) Cortical neurons expressing MitoDsRed (green) to label mitochondria (as well as GCaMP6s to measure neuronal activity, see Figure S4) fixed and immunostained with anti-O-GlcNAc (RL2, fire LUT) antibody immediately after the field stimulation (10Hz, 600AP) or under baseline conditions. (F) The total intensity of O-GlcNAc immunofluorescence at the dendritic and axonal mitochondrial compartments quantified from non-stimulated or stimulated neurons. Data are shown as Min-Max Box-whisker plot with associated p-values (n.s.= p>0.05) (Mann-Whitney U test). n = 646-801 mitochondria from 17-21 neurons from three independent experiments. Scale bar = 5 µm. See also Figure S4.

Given that OGT activity is regulated by glucose flux and the availability of its substrate UDP-GlcNAc (Figure 4A) ^8^, we hypothesized that mitochondrial O-GlcNAcylation is regulated by activity-dependent glucose metabolism. We therefore examined O-GlcNAcylation in neurons expressing the mitochondrial marker Mito-DsRed immediately following acute neuronal stimulation. An immunocytochemical analysis revealed that mitochondrial O-GlcNAcylation was enhanced in neuronal processes after the induction of neuronal activity (Figure 4E–F). Because O-GlcNAcylation is a dynamic post-translational modification, we blocked the removal of O-GlcNAc from proteins throughout the electrical stimulation by inhibiting the O–GlcNAc-removing enzyme O-GlcNAcase (OGA) using Thiamet-G (Figure S4D). This acute inhibition of OGA increased the accumulation of mitochondrial O-GlcNAcylation during baseline neuronal activity (Figure S4D–E), and this accumulation became more prominent upon neuronal stimulation. These results indicate that mitochondrial O-GlcNAcylation is a dynamic process in neurons that is rapidly promoted by neuronal activity and activity-dependent glucose metabolism.

### O-GlcNAcylation regulates mitochondrial respiration in neurons

We next assessed how O-GlcNAcylation regulates mitochondrial bioenergetics in neurons. We blocked the removal of O-GlcNAcylation by treating neuronal cultures with the OGA inhibitor Thiamet-G overnight and verified that this treatment increased O-GlcNAcylation (Figure S5). Then, we evaluated mitochondrial activity by double-labeling neurons with the mitochondrial membrane potential (ΔΨ_m_)-sensitive fluorescent dye tetramethylrhodamine methyl ester (TMRM) and the ΔΨ_m_-insensitive mitochondrial probe MitoTracker Green (MT Green) as a control. At sub-quenching concentrations, TMRM rapidly equilibrates in mitochondria and reflects ΔΨ_m_. Thiamet-G treatment increased ΔΨ_m_, which is quantified as the ratio between the intensities of TMRM and MT Green (Figure 5A–B). To evaluate mitochondrial respiratory capacity, we measured the mitochondrial oxygen consumption rate in control and Thiamet-G-treated neurons. As shown with neuronal activity (Figure 3H–K), changes in mitochondrial respiration did not seem to drive basal mitochondrial ATP synthesis, though the maximal respiratory capacity was enhanced in Thiamet-G-treated neurons (Figure 5C–E). We confirmed that the mitochondrial activity boost was not due to mitochondrial biogenesis, as Thiamet-G-treated neuronal mitochondrial protein levels remained constant (Figure S5). Thus, our results suggest that O-GlcNAcylation plays an important role in regulating ΔΨ_m_ and promotes mitochondrial respiratory capacity in neurons.

**Figure 5.**
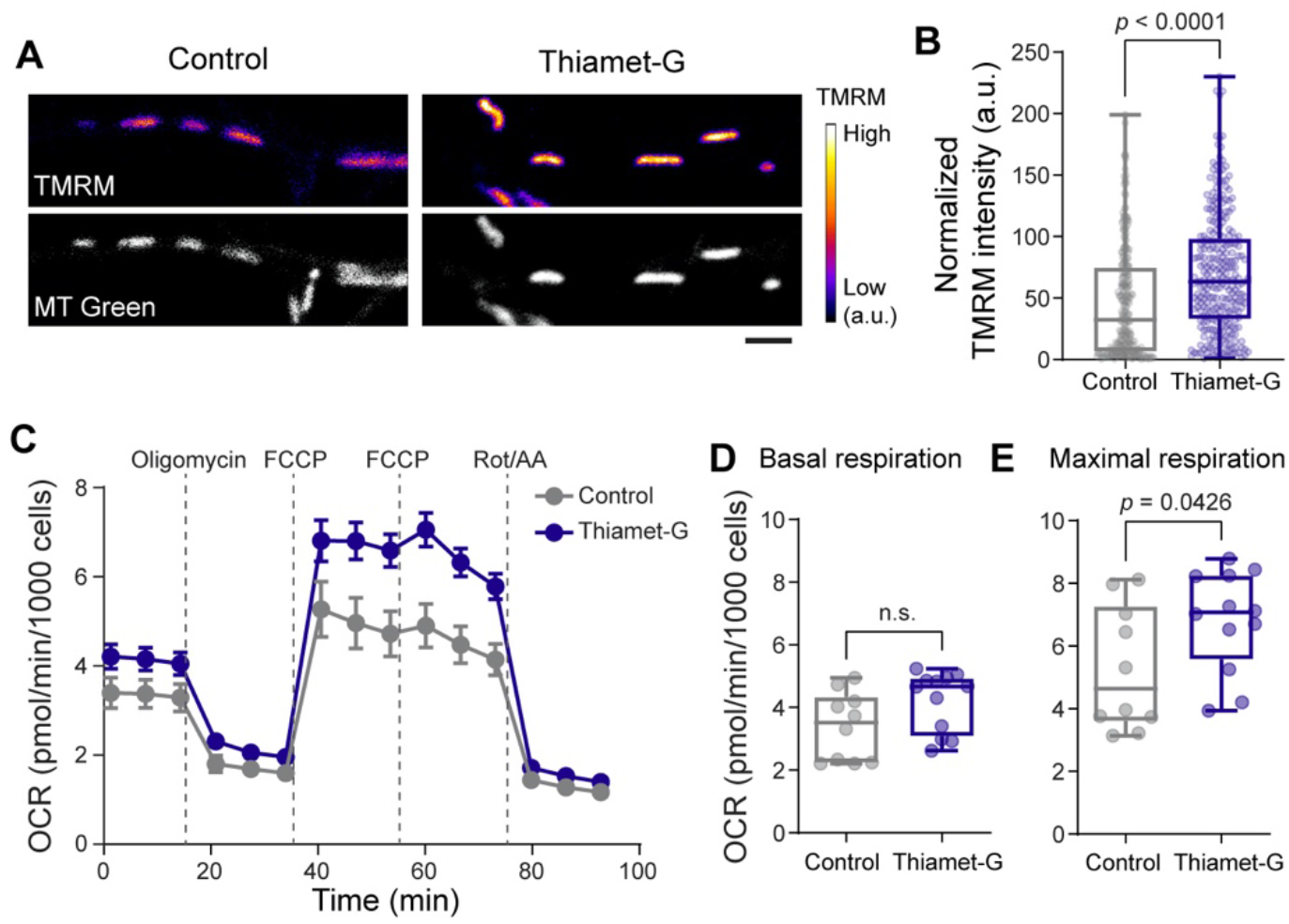
O-GlcNAcylation enhances mitochondrial activity. (A) Representative images demonstrating co-labeling of total mitochondria with MitoTracker (MT) Green (gray) and mitochondrial membrane potential with TMRM (fire LUT). (B) Mitochondrial membrane potential changes are represented as “Normalized TMRM intensity” by calculating the ratio of TMRM and MT Green intensities for each mitochondrion from neuronal processes. n= 268-281 mitochondria, four independent experiments. Data are shown as a Min-Max Box-whisker plot with associated p-values (Mann-Whitney U test). Scale bar = 2 µm. (C) Mitochondrial oxygen consumption rate (OCR) was measured from cortical neuron cultures using the “mito stress test” protocol after treatments with DMSO (vehicle control) or Thiamet-G overnight. (D-E) Basal (OCR_basal_) and maximal respiration (OCR_FCCP_) for indicated treatments. n= 10-12 wells per condition from three independent experiments. Data are shown as a Min-Max Box-whisker plot with associated p-values (n.s.= p>0.05) (Mann-Whitney U test). See also Figure S5.

### Mitochondrial O-GlcNAcome reveals adaptive bioenergetic mechanisms

Only a fraction of O-GlcNAcylated proteins undergo dynamic and reversible modification ^24^. To specifically identify the mitochondrial proteins that are O-GlcNAc modification when maximal respiratory capacity is enhanced (Figure 5), we developed a workflow for the comparative O-GlcNAcome profiling of neuronal mitochondria via quantitative mass spectrometry analysis. The mitochondria were purified from primary neuron cultures treated with Thiamet-G or vehicle control. We prepared crude-mitochondrial fractions, as opposed to ultra-pure mitochondria, and minimized the sample preparation steps/duration to preserve the chemically labile O-GlcNAc modification. We then treated mitochondrial lysate with succinylated wheat germ agglutinin (sWGA) agarose beads to selectively enrich for O-GlcNAcylated proteins (Figure 6A). We confirmed the binding specificity of O-GlcNAc-modified proteins to these beads using negative controls in the presence of free GlcNAc. sWGA-bound fractions obtained from Thiamet–G-treated neurons displayed a higher level of O-GlcNAcylated proteins, confirming our approach to capture the mitochondrial proteins that undergo dynamic O-GlcNAcylation (Figure S6A–B).

**Figure 6.**
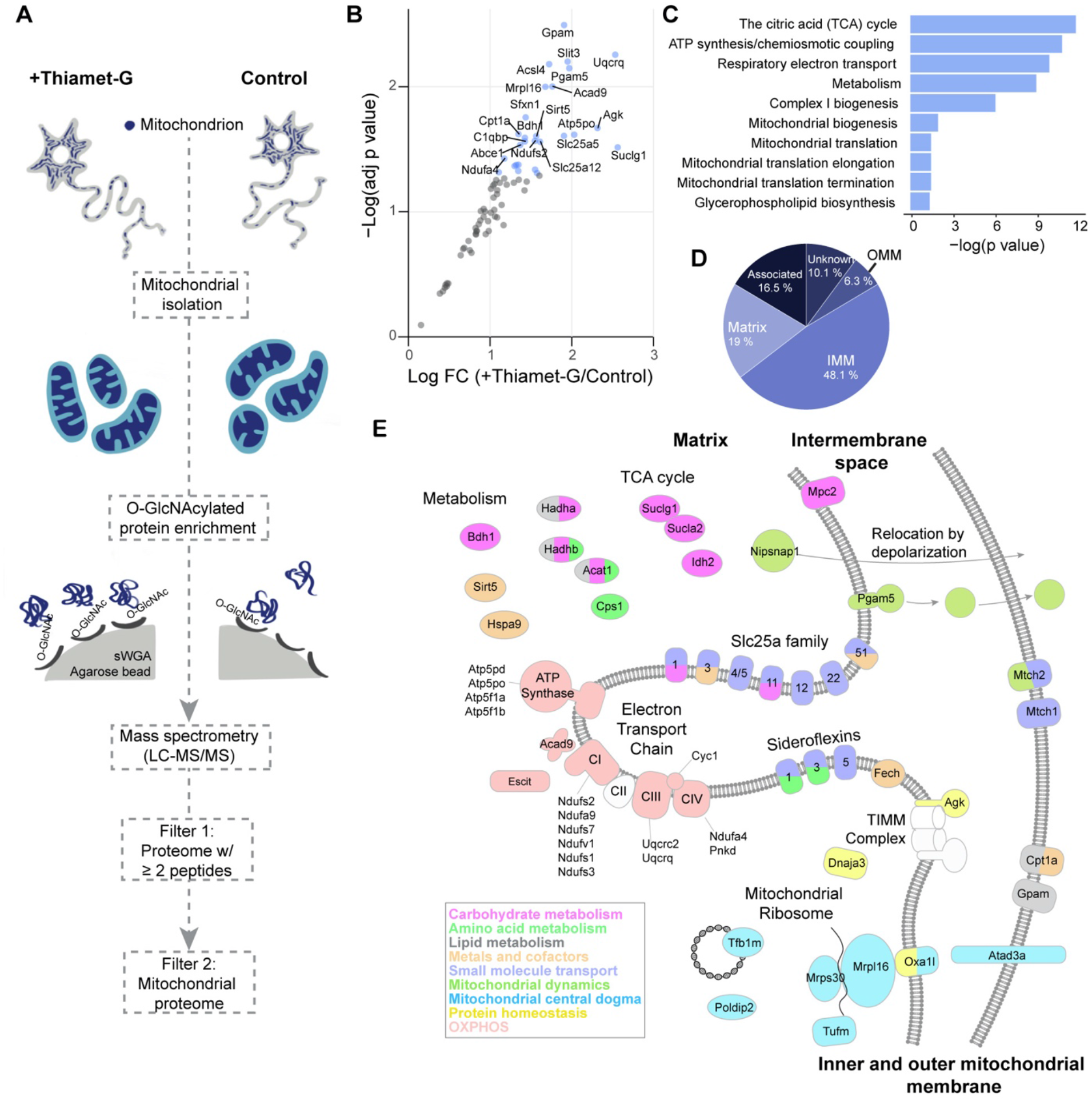
Mitochondrial O-GlcNAcome reveals mitochondrial plasticity mechanisms. (A) Schematic outline of mass spectrometry sample preparation and analysis strategies. Cortical neurons at 12-15 DIV were treated with DMSO (vehicle control) or Thiamet-G overnight to capture dynamic O-GlcNAc modification. O-GlcNAc-modified proteins were enriched from crude mitochondrial fractions using succinylated wheat germ agglutinin (sWGA) beads. Label-free quantitative mass spectrometry (LC-MS/MS) data was further analyzed to select proteins with ;: 2 peptides (Filter 1) and to identify mitochondrial proteins based on MitoCarta 3.0 and Mitominer databases (Filter 2). (B) Scatter plot analysis demonstrating the impact of Thiamet-G treatment on O-GlcNAcylated mitochondrial proteins. Annotated proteins (blue dots) represent p<0.05 (FC, fold change). (C) Pathway enrichment analysis, and (D) sub-mitochondrial localization of identified mitochondrial proteins. (E) Illustration of mitochondrial O-GlcNAcome, colored by MitoPathway assignments from MitoCarta3.0. n=3 biological replicates. See also Figure S5, Table S1 and S2.

To maximize the stringency in our subsequent mass spectrometry analysis, we selected proteins with ≥ 2 unique peptides in all experimental replicates from Thiamet-G or vehicle-treated samples. Using these selection criteria, we identified a total of 770 proteins (Table S1 and S2), 80 of which (∼10%) had confirmed mitochondrial localization based on the Mitominer and Mitocarta databases ^25,26^. The remaining were non-mitochondrial proteins, mostly localized in the cytosol (23.3%), plasma membrane (10.9%), or endoplasmic reticulum (ER) (10.8%) (Figure S6C–D), which was expected given the use of crude mitochondrial fractions. According to the O-GlcNAcome catalog ^27^, 142 of these proteins were previously identified to be O-GlcNAc modified (Figure S6C). As predicted, treatment with Thiamet-G significantly increased O-GlcNAcylation of mitochondrial proteins in neurons (Figure 6B–C and S6E).

We next analyzed the sub-mitochondrial localization of O-GlcNAcylated mitochondrial proteins (Figure 6C–E), finding that the majority were associated with the mitochondrial inner membrane (MIM) (48.1%) and matrix (19%), with only a minor proportion coming from the outer mitochondrial membrane (OMM) (Figure 6D). This further confirmed the involvement of this post-translational modification in regulating mitochondrial bioenergetics. Finally, we examined the specific mitochondrial functions regulated by O–GlcNAc-modified proteins using a protein-protein interaction network and pathway enrichment analyses. The top hits here were proteins involved in the tricarboxylic acid (TCA) cycle, ATP synthesis, electron transport chain function, and metabolite transport (Figure 6B–E, S6E and Table S1-2). Thus, O-GlcNAc modification of mitochondrial proteins may support metabolic plasticity in neurons by coordinating multiple pathways involved in energy metabolism.

### O-GlcNAc transferase supports on-demand ATP synthesis

Because the majority of O-GlcNAcylated mitochondrial proteins enriched in our proteomics data (Figure 6B–C) are involved in pathways supporting ATP synthesis, we hypothesized that O-GlcNAcylation is involved in neuronal activity-driven on-demand ATP synthesis. To test our hypothesis, we expressed the cytoplasmic ATP sensor iATPSnFR1.0-mRuby in neurons and triggered AP firing (10 Hz, 600 AP) in the presence of the vehicle control or OGT inhibitor OSMI-4 (Figure 7A and S7A–B). By imaging intracellular Ca^2+^ with GCaMP6s before and after neuronal stimulation, we confirmed that the application of OSMI-4 (or OGA inhibitor Thiamet-G) to prevent the addition of O-GlcNAc did not alter neuronal activity (Figure S7C–E). Measurements from neuronal processes in control cells revealed that a burst of electrical activity resulted in only a small decrease in ATP levels during the post-stimulus period, which was followed by an immediate recovery (Figure 7A). On the contrary in OSMI-4-treated neurons, ATP levels never recovered and instead decreased by ∼30% from the baseline at 10 min post-stimulation (Figure 7B). When both glycolytic and mitochondrial (2DG + Oligo) ATP synthesis pathways were blocked during electrical stimulation, the ATP level decreased >60% in neurons. This result, combined with the observation that neuronal activity enhances glucose metabolism as well as mitochondrial O-GlcNAcylation (Figure 4), implies that OGT activity is required for the upregulation of neuronal bioenergetics.

**Figure 7.**
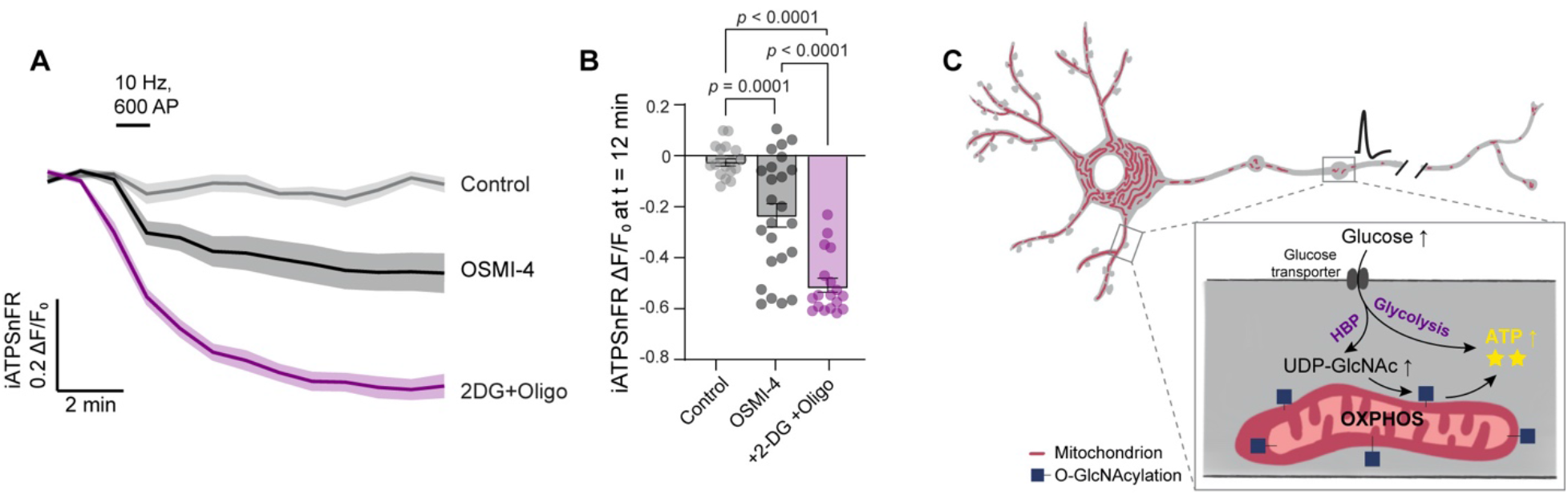
O-GlcNAc transferase supports on-demand ATP synthesis. (A) Average fluorescence traces of ATP sensor, iATPSnFR, with indicated pharmacological treatments and field stimulation (10 Hz, 600AP) (gray: control; black: OGT inhibitor OSMI4; magenta: 2DG and Oligomycin). Data are shown as mean values ± SEM. (B) Quantification of ATP depletion (iATPSnFR ι1F/F_0_) from neuronal processes at 10 minutes post-stimulation (t = 12 min) for each condition. Data are shown as and Min-Max Box-whisker plot with associated p-values (one-way ANOVA with post hoc Kruskal-Wallis multiple comparison test), n = 14-24 neuronal processes, 4-8 cells, and three independent experiments. (C) Summary diagram demonstrating neuronal mitochondria (red) and glucose metabolism (magnified image with dashed box). Neuronal activity stimulates glycolysis and UDP-GlcNAc synthesis via the hexosamine biosynthetic pathway (HBP), which then enhances O-GlcNAcylation (blue squares) of mitochondrial proteins for “on-demand” ATP synthesis. See also Figure S7.

## DISCUSSION

Metabolic flux-sensitive post-translational modification, O-GlcNAcylation, uniquely couples fuel availability to cellular metabolism and signaling pathways ^28^. Here, we demonstrate how O-GlcNAc modification magnifies the functional diversity of mitochondrial proteins and functions in neurons. We have now established that (1) neuronal activity drives fuel consumption and consequently dynamic O-GlcNAcylation in neuronal processes; (2) especially mitochondria are the loci of O-GlcNAc modification; (3) post-translational modification of mitochondrial proteins via OGT enhances mitochondrial respiratory capacity in neurons; and (4) mitochondrial O-GlcNAcome and O-GlcNAc cycling supports neuronal activity-driven on-demand ATP synthesis and energy homeostasis.

O-GlcNAcylation level in neurons correlated well with neuronal activity, when the activity was driven by chronic pharmacological manipulations *in vivo* (Figure 1) and *in vitro* (Figure 2) or by acute electrical stimulation (Figure 4). O-GlcNAcylation is highly enriched in the central nervous system^29,30^ and essential for brain function ^24,31-33^. While previous biochemical studies indicated that neuronal activity enhances O-GlcNAcylation in the brain ^24^, whether non-neuronal cells or neurons are the primary target of this post-translational modification was ambiguous. By using immunofluorescence staining and cellular analysis, we established that neurons, specifically neuronal processes and mitochondria are the loci of this modification. OGT expression and O-GlcNAcylation is developmentally regulated in neurons ^31,33,34^. We demonstrated that in mature neurons, O-GlcNAcylation was enriched in neuronal processes (Figure S2 and 2). Increase in axonal and dendritic O-GlcNAcylation also coincides with enhanced energy demand, synaptogenesis, and metabolic shift from glycolysis to OXPHOS ^35,36^. This supports the critical role of O-GlcNAcylation in sustaining metabolic homeostasis by coupling enhanced energy demand and associated fuel consumption in neuronal processes.

Glucose is the major fuel source of the neurons. In response to increased energy demand, neuronal glucose consumption and glycolysis can be upregulated ^4-6^. Long-term increase in glucose flux (1-2 hours), also leads to increased O-GlcNAcylation in neurons ^8^. However, we show that mitochondrial O-GlcNAcylation is rapidly upregulated within minutes upon activity-driven glucose utilization (Figure 4). Neuronal glucose flux has been shown to immobilize mitochondria at high-glucose areas by increasing the O-GlcNAc modification of motor-adaptor protein Milton and anchoring mitochondria to actin cytoskeleton ^8,13^. We find that global enhancement of O-GlcNAcylation by the pharmacologically inhibition OGA, a manipulation that arrests mitochondrial motility ^8,13^, also enhances mitochondrial O-GlcNAcylation and promotes mitochondrial respiratory capacity in neurons. Co-regulation of mitochondrial localization and activity in fuel-rich areas allows neuronal mitochondria to rapidly adapt local metabolic demand to sustain neuronal energy homeostasis. We specifically investigated which mitochondrial proteins are dynamically regulated via O-GlcNAcylation and identified proteins involved in ATP synthesis that include electron transport chain subunits, ADP phosphate carriers, malate-aspartate shuttle subunits (Figure 6). Because we worked with crude mitochondrial fractions, our O-GlcNAcome data uncovered other neuronal proteins including pre-synaptic proteins, endo-lysosomal membrane proteins, and cytoskeletal components (Figure S6). These results may suggest that neuronal activity-driven O-GlcNAcylation may be a potential way to synchronize multi-organelle functions based on fuel availability in neurons.

The origin of mitochondrial O-GlcNAcylation is controversial ^37-39^. A single pair of enzymes, OGA and OGT, regulates nuclear, cytoplasmic and mitochondrial protein O-GlcNAcylation in cells. Although both enzymes are encoded by a single gene, their splice isoforms are identified in different cellular compartments including mitochondria ^10,37,40^. It is possible that mitochondrial proteins are O-GlcNAcylated in the cytoplasm, then subsequently imported into mitochondria, however, inner membrane localization of mitochondrial OGT^37,40^ and OGA^41^ as well as the presence of mitochondrial UDP-GlcNAc transporter^40^ supports O-GlcNAcylation occurring in mitochondria ^42^. The spatiotemporal regulation of O-GlcNAc cycling is complex and regulated via multiple different mechanisms. While fuel availability can lead to global changes in O-GlcNAc level, binding partners of the regulatory enzymes OGA and OGT can fine-tune O-GlcNAcylated substrates locally in different subcellular compartments. Future work is required to identify the role of each OGT/OGA splice isoforms and GlcNAc modification in the regulation of mitochondrial plasticity in neurons.

Mitochondrial respiration is mainly controlled by fuel availability and the ATP turnover rate ^43^. In neurons, electrical activity consumes ATP and enhances fuel utilization (Figure 4) for instantaneous ATP synthesis ^3,6^. Mitochondrial Ca^2+^ uptake is proposed as the critical regulator of activity-driven mitochondrial ATP synthesis, though downstream mechanisms have not been explored in neurons ^44-47^. We report a new mechanism that uses post-translational modification, O-GlcNAcylation, to couple neuronal activity, metabolism and mitochondria to support on-demand ATP synthesis. Mitochondrial O-GlcNAcylation is rapidly upregulated by activity-dependent glucose utilization (Figure 4). Both calcium and glucose may stimulate on-demand ATP synthesis by providing feedforward regulation. While mitochondrial Ca^2+^ uptake may initiate instantaneous on-demand mitochondrial activity, associated with Ca^2+^ entry from the plasma membrane, O-GlcNAcylation may provide continuous tailoring of mitochondrial ATP generation based on fuel availability. We identified proteins involved in the ketone body metabolism enzymes, i.e., D-beta-hydroxybutyrate dehydrogenase (BDH1) and metabolite transporters in our mitochondrial O-GlcNAcome (Figure 6). Although glucose is the main fuel source in the brain, neurons have the ability to adapt to changes in nutrient availability and utilize different energy substrates ^48,49^. Conceivably, nutrient sensing via the HBP pathway and OGT activity inform mitochondria about the source of fuel and allow mitochondrial metabolism to be tailored based on nutrient availability.

Mitochondrial function and neuronal O-GlcNAcylation impairments are associated with neurodegenerative disorders including Alzheimer’s, Huntington and Parkinson’s disease ^1,50-54^. Upregulating of O-GlcNAc modification protects against neurodegeneration, motor deficits and synaptic impairment; however, biological stress, such as epileptiform activity, and aging deplete brain O-GlcNAcylation ^55-58^. Compromised glucose metabolism and metabolic defects are also the main contributors of neurodegenerative diseases. Our study demonstrates a new link between neuronal glucose metabolism, O-GlcNAc modification and mitochondrial bioenergetics, thus providing important insight on the metabolic basis of neurodegeneration.

## ACKNOWLEDGEMENTS

We thank members of the G.P. laboratory, Dr. Ajit Divakaruni, Dr. Jeffrey Esko, Dr. Gerald Shadel, Dr. Anthony Molina, Dr. Lance Wells, Dr. Matthew Banghart, and Dr. Susan Ackerman for their valuable suggestions. We are grateful to Dr. Richard Goodman and Dr. Philip J. S. Stork for the support and guidance with the HYlight sensor experiments. We also thank the members of the University of California San Diego BPMSF and Center for Computational Biology and Bioinformatics Core for their technical contributions. This work was supported by a grant from NIH (R35GM128823) to G.P., NIH (2T32GM007240-39A1) to S.B.Y., University of California San Diego David V. Goeddel endowed graduate fellowship to S.B.Y. and San Diego IRACDA Scholars Program (K12GM068524) grant to R.S.

## AUTHOR CONTRIBUTIONS

G.P. and S.B.Y. designed the study. S.B.Y., R.S., and Z.D.P. performed the experiments. J.N.K. and M.L.S. provided the HYLight sensor and feedback on the experimental design. M.G. and T.C.W. carried out the mass spectrometry experiments and analysis. G.P. and S.B.Y. wrote the manuscripts with input from co-authors.

## DECLARATIONS OF INTERESTS

The authors declare no competing interests.

## FIGURE LEGENDS

**Figure S1.**
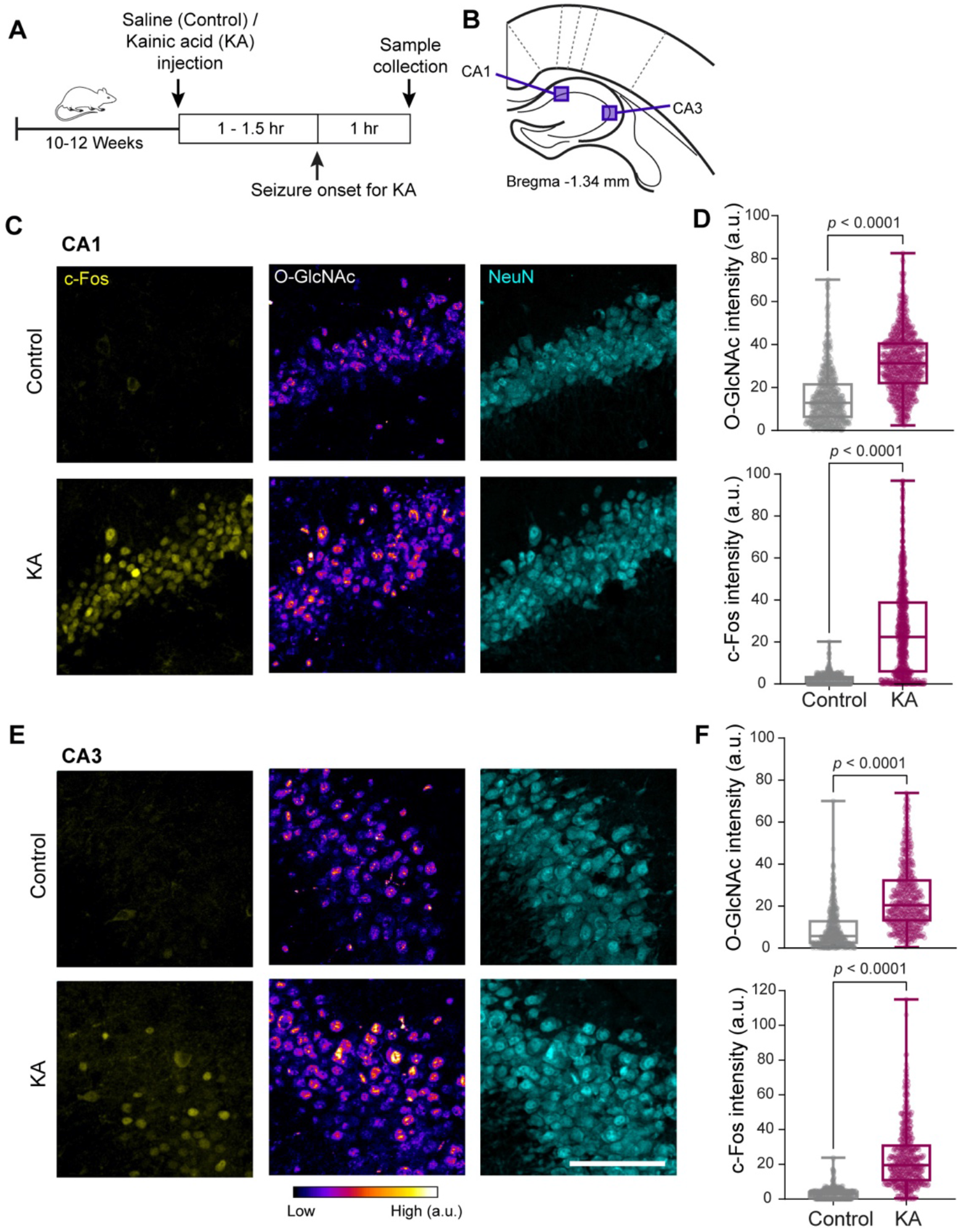
O-GlcNAcylation increases with neuronal activity in the CA1 and CA3 regions of the hippocampus. Related to Figure 1. (A) Scheme demonstrating the experimental timeline of intraperitoneal saline or kainic acid injections, seizure induction, and sample collection. (B) O-GlcNAc and c-Fos fluorescence intensities were analyzed in the indicated hippocampal areas (CA1 and CA3) from brain sections ranging from Bregma -1.22 to -1.45 mm. (C-F) Representative images of CA1 and CA3 hippocampal regions after saline (Control) or kainic acid (KA) injections and immunohistochemical staining with antibodies against c-Fos (yellow), O-GlcNAc (RL2, fire LUT), and NeuN (cyan). (D and F) Quantification of c-Fos and O-GlcNAc intensities from NeuN-positive neurons in (D) CA1 and (F) CA3 hippocampal regions from saline (grey) and KA injected (magenta) brains. For each condition n = 655-811 neurons in CA1 and 508-566 neurons in CA3, 3 mice. Data are shown as a Min-Max Box-whisker plot with associated p-values (Mann-Whitney U test, Scale bars = 100 µm).

**Figure S2.**
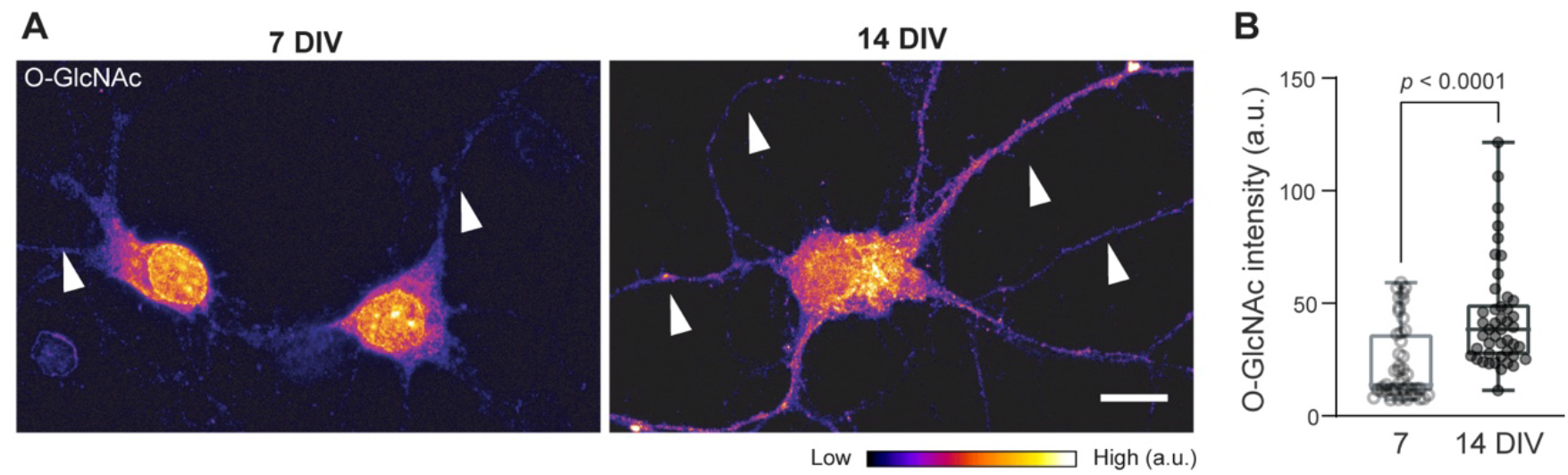
Changes in neuronal O-GlcNAcylation level in primary neuron cultures. Related to Figure 2. (A) Representative images of 7 and 14 days *in vitro* (7 or 14DIV) primary cultured cortical neurons immunostained with anti-O-GlcNAc (RL2, fire LUT) antibody to visualize O-GlcNAcylation levels. The arrowheads indicate the enrichment of O-GlcNAcylation in neuronal processes as neurons mature. Scale bar = 10 µm. (B) Quantification of O-GlcNAc fluorescence intensities from 7 and 14 DIV neurons. Data are shown as a Min-Max Box-whisker plot with associated p-values (Mann-Whitney U test). n= 45 cells from three independent experiments.

**Figure S3.**
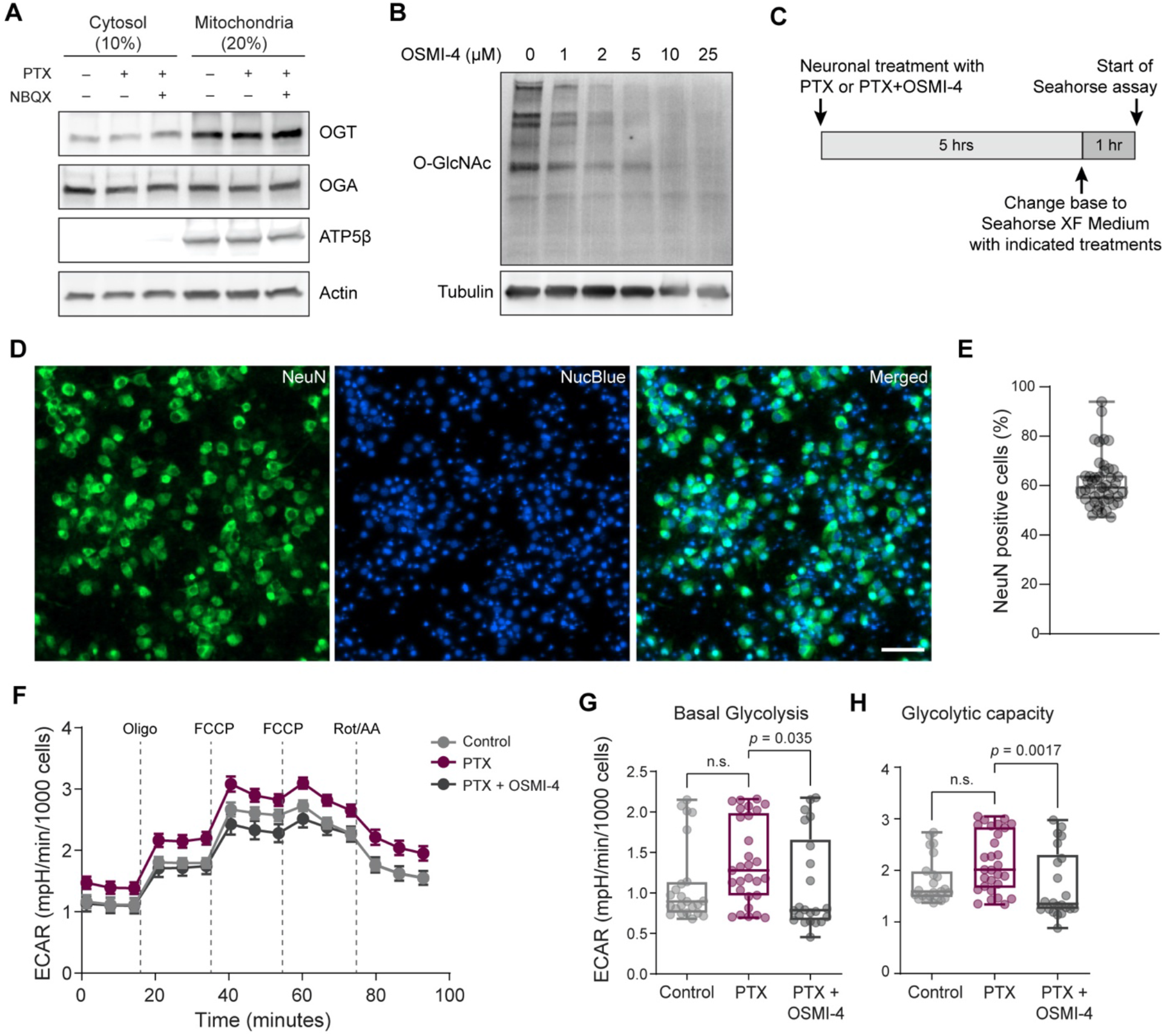
Characterization of cortical neuron cultures and O-GlcNAc manipulations. Related to Figure 3. (A) OGT and OGA protein levels from mitochondrial and cytosolic fractions, obtained from cortical neurons treated with DMSO (vehicle control), PTX, or PTX+NBQX for 6 hours, analyzed by SDS gel electrophoresis and probed with anti-OGT, anti-OGA, anti-ATP5β and anti-actin antibodies. (B) The total protein O-GlcNAcylation level changes were evaluated after different concentrations of OGT inhibitor OSMI-4 treatments (0, 1, 2, 5, 10, and 25 µM) for 6 hours from cortical neuron lysates. Cell lysates were separated by SDS gel electrophoresis and probed with anti-O-GlcNAc antibodies (RL2) and anti-tubulin (loading control) antibodies. (C) Schematic of the experimental regimen for respirometry measurements with Agilent Seahorse XFe96 Analyzer. (D) Retrospective immunostaining of 96-wells plate with anti-NeuN (green) antibody and NucBlue (blue, nuclear marker) after respirometry measurements to examine the extent of neuronal enrichment in cortical cultures per well. (E) Percentage of NeuN positive cells calculated for each well. Data are shown as a Min-Max Box-whisker plot. n= 46 wells from three independent experiments. Scale bar = 50 µm. (F) Extracellular acidification rate (ECAR) per 1000 cells measured from cultured cortical neuron cultures using the “mito stress test” protocol after treatments with DMSO (vehicle control), PTX, or PTX+OSMI-4 for 6 hours. (I-K) Basal (ECAR_basal_) and glycolytic capacity (ECAR_Oligomycin_ - ECAR_basal_) for indicated treatments. Mean ± SEM for each time point, n= 24-32 wells per condition from three independent experiments. Data are shown as a Min-Max Box-whisker plot with associated p-values (n.s.= p>0.05) (one-way ANOVA with post hoc Tukey’s multiple comparison test).

**Figure S4.**
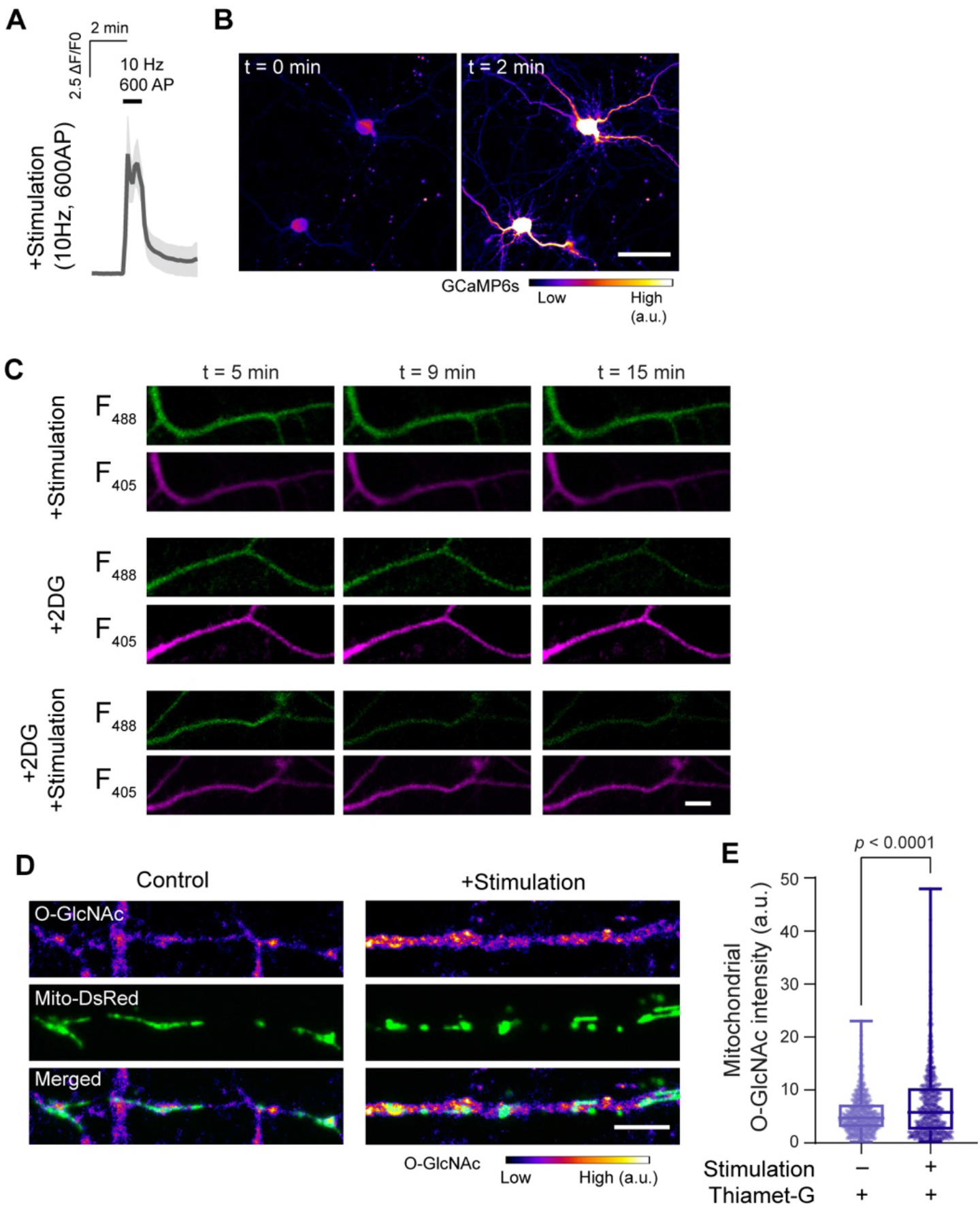
Activity-dependent calcium, glycolysis and O-GlcNAcylation measurement details. Related to Figure 4. (A-B) Change in somatodendritic fluorescence of GCaMP6s in response to a train of stimulation (10Hz, 600AP). (A) Average fluorescence traces of GCaMP6s (ΔF/F_0_) signals to demonstrate the cytosolic Ca^2+^ responses to electrical field stimulation. Data are shown as mean values ± SEM. n= 5-9 neurons from three independent experiments. (B) Representative image of the change in fluorescence of GCaMP6s (fire LUT) before (t = 0min) and after (t = 2min) electrical stimulation. Scale bar = 50 µm. (C) Representative images of neuronal processes expressing HYlight sensor, before (t= 5 min), and after (t= 9 and 15 min) field stimulation (10Hz, 600AP) either with or without 2DG treatments. The normalized HYlight emission ratio (demonstrated in Figure 4B) is induced by 488nm (F_488_) and 405nm (F_405_) excitations at indicated time points. Scale bar = 10 µm. (D) Cortical neurons expressing MitoDsRed (green) to label mitochondria (as well as GCaMP6s to measure neuronal activity) fixed and immunostained with anti-O-GlcNAc (RL2, fire LUT) antibody immediately after the field stimulation (10Hz, 600AP) or under baseline conditions. To block the removal of O-GlcNAc modification, neurons were treated with Thiamet-G one hour before the initiation of imaging and electrical stimulation. (E) The total intensity of O-GlcNAc immunofluorescence at the dendritic and axonal mitochondrial compartments quantified from non-stimulated or stimulated neurons. Data are shown as a Min-Max Box-whisker plot with associated p-values (n.s.= p>0.05) (Mann-Whitney U test). n = 568-634 mitochondria from 19-23 neurons, three independent experiments. Scale bar = 5 µm.

**Figure S5.**
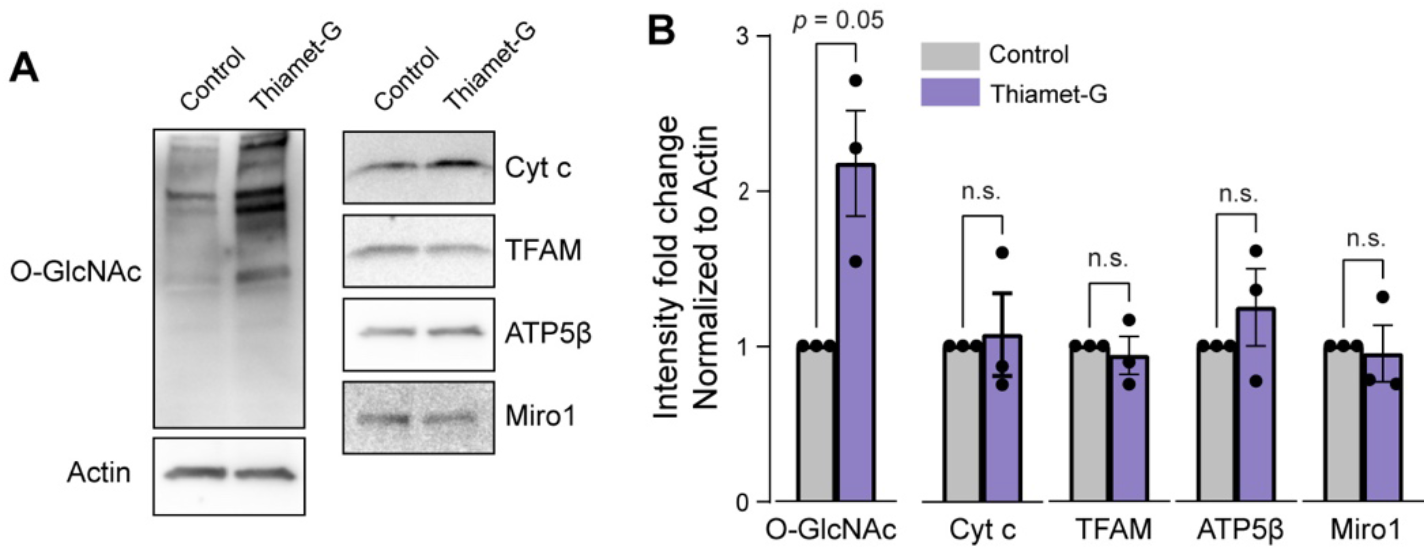
Evaluation of neuronal mitochondrial content. Related to Figure 5. (A) The total protein O-GlcNAcylation level and mitochondrial protein content changes were evaluated from cortical neuron lysates, after overnight vehicle or Thiamet-G treatments. Cell lysates were separated by SDS gel electrophoresis and probed with anti-O-GlcNAc (RL2), anti-actin (loading control) and mitochondria-specific anti-cytochrome c (Cyt c), anti-TFAM, anti-ATP5β, anti-Miro1 antibodies. (B) The total intensity of the O-GlcNAc, Cyt c, TFAM, ATP5β and Miro1 immunoreactive bands were normalized to the intensity of actin bands for each lane. Fold changes in response to indicated treatments were calculated. n = 3 independent experiments. Data are shown as mean values ± SEM with associated p-values (Mann-Whitney U test).

**Figure S6.**
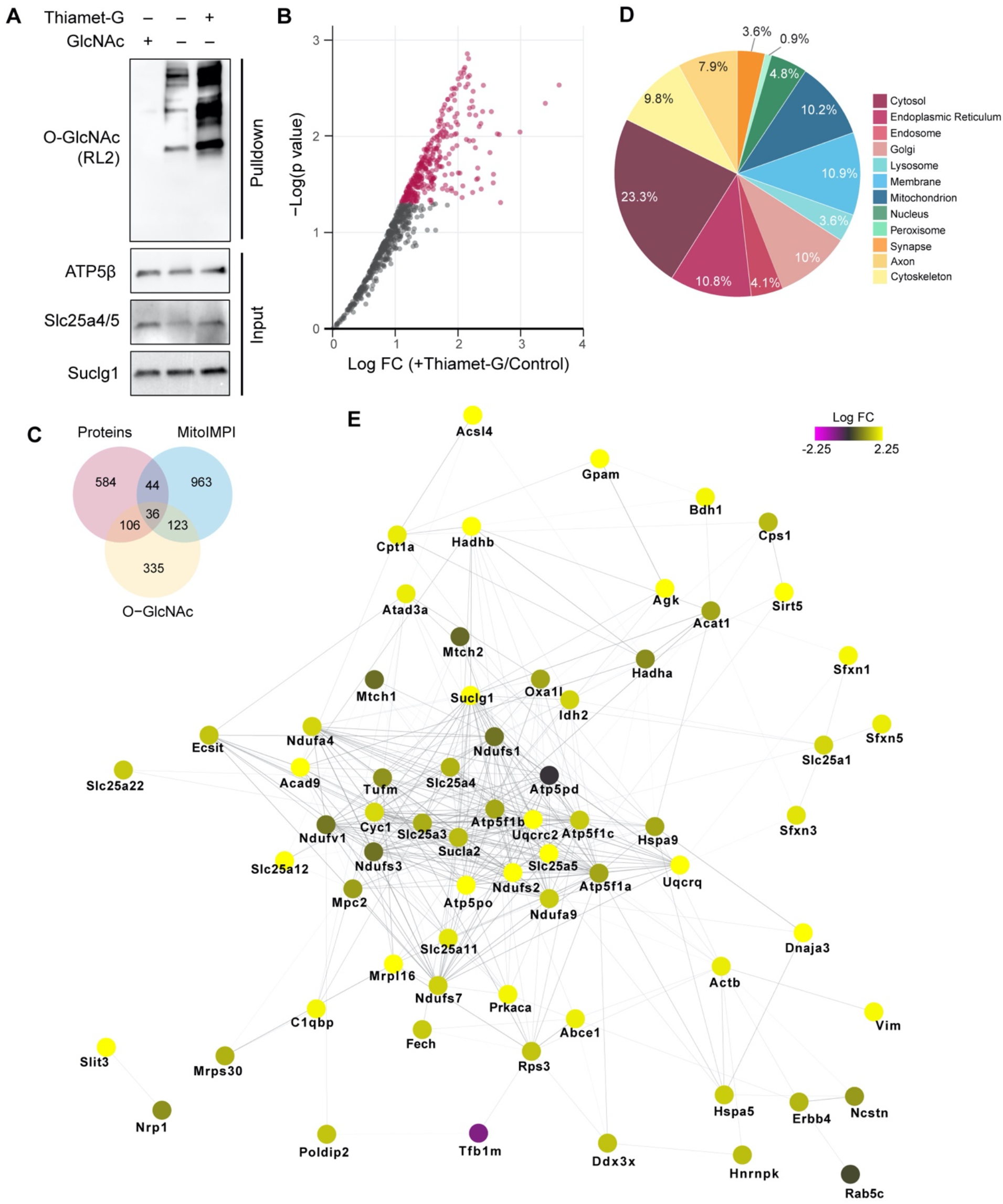
Subcellular analysis of O-GlcNAcylated proteins. Related to Figure 6. (A) The total sWGA-bound protein O-GlcNAcylation level changes evaluated by SDS gel electrophoresis and probed with anti-O-GlcNAc antibodies (RL2). Crude mitochondrial fractions isolated from cortical neurons at 12-15 DIV, treated with DMSO (vehicle control) or Thiamet-G overnight to augment O-GlcNAc modification. O-GlcNAc modified proteins were enriched from crude mitochondrial fractions using succinylated wheat germ agglutinin (sWGA) beads. Free GlcNAc used as a control to validate the specificity of sWGA binding. Mitochondrial fractions (input) were analyzed by SDS gel electrophoresis and probed with anti-ATP5β, anti-Slc25a4/5 and anti-Suclg1 antibodies. and anti-tubulin (loading control) antibodies. (B) Scatter plot analysis demonstrating the impact of Thiamet-G treatment on O-GlcNAcylated proteins. Annotated proteins (red dots) represent p<0.05 (FC, fold change). (C) Venn diagram analysis of identified proteins using mitochondrial databases (MitoCarta3.0 and IMPI) and O-GlcNAcome catalogue. (D) Subcellular localization of identified proteins. (E) STRING protein-protein interaction analysis of mitochondrial proteins with nodes colored by FC Thiamet-G / Control.

**Figure S7.**
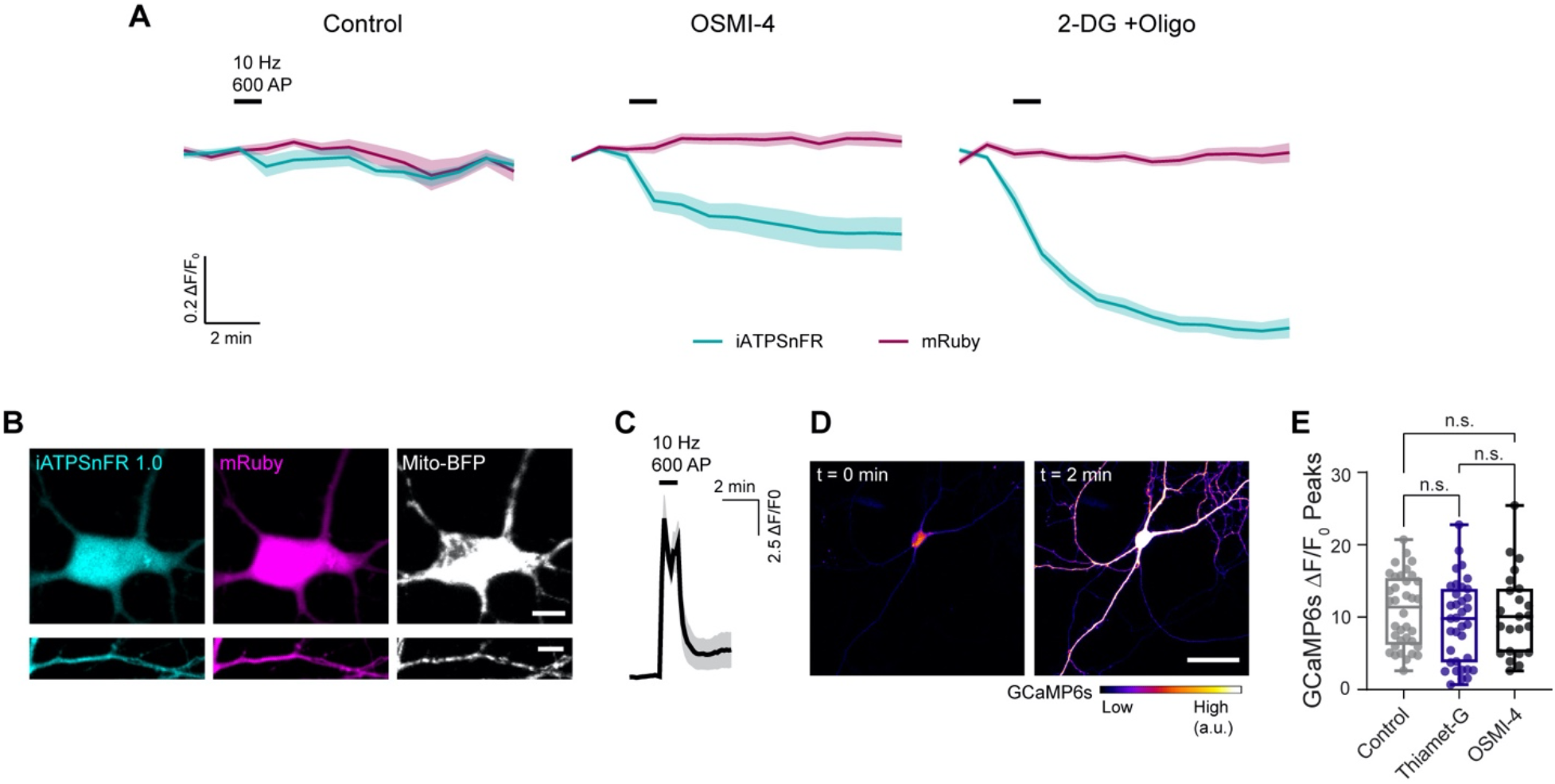
Acute alterations of ATP and Calcium levels in response to OGT inhibitor treatment. Related to Figure 7. (A) Traces of iATPSnFR and mRuby F/F0 over time for each condition, 2 minutes before, during, and 10 minutes after the field stimulation (10 Hz, 600 AP). (B) Images of cortical neuronal soma and processes transfected with iATPSnFR1.0-mRuby and Mito-BFP. (C) Normalized GCaMP6s ΔF/F_0_ signal from stimulated neurons, pre-treated with 5 μm OSMI-4 for 30 min. The trace represents mean values ± SEM from 5-9 neurons from three independent experiments. (D) Pseudocolored representative neuronal GCaMP6s before (t = 0 min) and after (t = 2 min) electrical stimulation. Scale bar = 50 µm. (E) Peak ΔF/F_0_ of GCaMP6s signals from individual neurons pre-treated with DMSO (Control), 5 μm OGA inhibitor Thiamet-G, and 5 μm OGT inhibitor OSMI-4, The peak values show no differences with pre-incubation of drugs. n = 23-36 ROIs taken from 6-9 neurons from three independent experiments.

## STAR METHODS

### RESOURCE AVAILABILITY

#### Lead contact

Further information and requests for resources should be directed and will be fulfilled by the lead contact, Gulcin Pekkurnaz.

## Materials availability

All biological resources and tools are either available from commercial sources or lead contact.

## Data and code availability

Proteomics data are provided in Supplementary Tables S1 and S2. The data analysis code is available at https://github.com/ucsd-ccbb/MitoProteomicsMethods. Agilent Seahorse XF96e Metabolic Flux Analyzer data normalization is performed via the custom-written macro “FluxNorm”, available at https://github.com/pekkurnazlab/FluxNormalyzer. Any additional information required to reanalyze the data reported in this paper is available from the lead contact upon request.

### EXPERIMENTAL MODEL AND SUBJECT DETAILS

#### Animals

All animal housing, breeding, and procedures were conducted according to the NIH Guide for the Care and Use of Experimental Animals and approved by the University of California San Diego Animal Care and Use Committee. C57BL/6J strain wild-type mice (Jackson Laboratory) were used for kainic acid and saline injections. Mice were housed at 22-24 °C in a room using a 12-hour light/dark cycle with *ad libitum* access to standard chow diet and water. Sprague-Dawley strain wild-type rats (Envigo/Harlan) were used for primary neuron cultures. Timed pregnant female rats (embryonic days 13-16) were singly housed in a room with a 12-hour light/dark cycle with *ad libitum* food, water access, and environmental enrichment.

#### Primary neuronal culture

Hippocampal and cortical neurons were isolated from rat (Envigo) embryos (E18) as previously described ^59^ and plated at 5-7 × 10^4^ cells/cm^2^ on coverslips for imaging or on 6-well plates at 1-2 × 10^5^ cells/cm^2^ for biochemical assays. Before plating, the No.1 12mm coverslips (Carolina Biological Supplies) and plates were coated with 20 µg/mL Poly-L-Lysine (Sigma-Aldrich) and 3.5 µg/mL Laminin (Life Technologies) overnight at room temperature. Primary neuron cultures were maintained in Neurobasal (NB) medium containing 5mM glucose, supplemented with B27, GlutaMAX, and penicillin/streptomycin (Life Technologies) unless modified as specified. Each independent experiment was performed by the preparation of new primary neuron cultures. Cortical neurons used for biochemical assays were treated with 1 µM Ara-C (Cytarabine, Tocris) at 3 days in vitro (DIV) to prevent glia proliferation for 2 days. Ara-C-containing medium was replaced with fresh neuron maintenance medium at 5 DIV. The primary neuron cultures were maintained for 11-15 days in vitro by replacing one-third of the culture medium with a fresh medium every three days.

## METHOD DETAILS

### Plasmid constructs

The following previously published or commercially DNA constructs were used: pDsRed2-Mito (Clontech Laboratories Inc.), Mito-BFP ^60^, HYLight ^23^. pAAV.CAG.GCaMP6s.WPRE.SV40 was a gift from Douglas Kim & GENIE Project (Addgene plasmid #100844; http://n2t.net/addgene:100844; RRID: Addgene_100844) ^61^. Synapsin-cyto-mRuby3-iATPSnFR1.0 was a gift from Baljit Khakh (Addgene plasmid # 102557; http://n2t.net/addgene:102557; RRID: Addgene_102557) ^62^.

### Kainic acid administration

Age-matched (10-12 weeks) C57BL/6J male mice were injected intraperitoneally with either 10 mg/kg kainic acid (KA) (5mg/ml solution using pharmaceutical grade saline) per dose or an equivalent volume of saline as a control. Kainic acid-injected mice were singly housed and monitored for visible signs of seizure including rearing and repetitive paw movements (Racine scale 4)^63^. If mice failed to reach Racine scale 4 seizure behavior within one-hour after KA injection, a booster dose (5 mg/kg) of KA was administered. One hour after the onset of Racine scale 4 seizure behavior, the paired saline and KA injected mice were anesthetized by inhalation of isoflurane and perfused transcardially first with ice-cold phosphate-buffered saline (PBS, pH 7.4) followed by freshly prepared 4% paraformaldehyde (PFA). The brain was extracted and postfixed with 4% PFA solution overnight, then rinsed with PBS, cryoprotected in 30% sucrose, and embedded with Tissue Tek OCT Compound for cryosectioning.

### Brain slice immunohistochemistry

Brains were sectioned at 25 µm on a cryostat (ThermoFisher Cryostart NX70). The sections were placed in blocking solution (5% Goat serum and 0.3% Triton X-100 in PBS) for 1 hour at room temperature, then incubated with primary antibodies in staining solution (1% Goat serum and 0.3% Triton X-100 in PBS) overnight at 4 ºC. After washing with PBS three times, secondary antibody staining was performed for 1 hour at room temperature. The sections were mounted using DAPI (4’,6-diamidino-phenyindole)-Fluoromount-G (Southern Biotech, Birmingham, AL). The following primary antibodies were used for staining: mouse anti-O-GlcNAc (RL2, Abcam, Ab2739), rabbit anti-c-Fos (Synaptic Systems 226003), chicken anti-NeuN (Sigma-Aldrich ABN91). The secondary antibodies were: goat anti-mouse Alexa 488 (Life Technologies A11029), goat anti-rabbit Alexa 568 (Life Technologies A11036), goat anti-chicken Alexa 647 (Life Technologies A21449), and DAPI for nuclear staining. Images were acquired using a Zeiss LSM780 confocal laser scanning microscope, Plan-Apochromat 20x/0.8 M27 objective using the same image acquisition settings across different conditions and sections. The images were further analyzed using Fiji ^64^, using only linear adjustments of brightness and contrast for visualization.

### Transfection and live-cell imaging

Primary cortical neuron cultures were transfected with indicated DNA constructs using Lipofectamine 2000 (Life Technologies). Live-cell imaging of neurons was performed 2-3 days after transfections at 11-12 DIV. Coverslips were mounted on a stimulation chamber (Warner Instruments RC-47FSLP) with laminar-flow perfusion and imaged at 37°C using Zeiss LSM780 laser scanning confocal microscope equipped with a heated stage and C-Apochromat 40x/1.20 W Korr FCS M27 objective. Laser power was set to <1% for each channel to minimize phototoxicity during time-lapse image acquisition. For most of the experiments, neurons were continuously perfused at 0.2-0.25 ml/min with Tyrode buffer (50mM HEPES pH 7.4, 119 mM NaCl, 2.5 mM KCl, 2 mM CaCl_2_, 2 mM MgCl_2_, 5 mM glucose, 2mM pyruvate, and 2mM lactate). Trains of action potentials were evoked by current pulses of 100mA, at 10Hz for 60 seconds during live-cell imaging to measure calcium dynamics, ATP, and glucose metabolism with genetically encoded sensors. The images were further analyzed as described below using Fiji ^64^.

### Calcium dynamics imaging and analysis

Neurons were transfected with fluorescent calcium sensor GCaMP6s to image neuronal activity as previously described ^61^. Time-lapse movies were acquired from 11-12 DIV neurons expressing GCaMP6s for a total of 5 minutes at 0.1-0.2 Hz. For each imaging session, first pre-stimulus fluorescence baseline (F_0_) measurements were performed for 2 minutes in the absence of electrical activity as a control, then action potentials were evoked by field stimulation (100mA,10 Hz,600 AP). GCaMP6s peak fluorescence (ΔF/F_0_) for each stimulation was calculated by dividing the highest intensity data point by pre-stimulus intensity. O-GlcNAc transferase and O-GlcNAcase activities were blocked by 5 µM OSMI-4 (MedChem Express) and 5 µM Thiamet-G (Calbiochem) treatments, respectively. Neurons were preincubated with OSMI-4 and Thiamet-G for 30 minutes before imaging and perfused with Tyrode buffer containing the inhibitors during live-cell imaging. For each neuron, at least three regions of interest (ROIs) with an average area of 20-30 µm^2^, encompassing all neuronal compartments (soma, dendrites, and axons) were selected for image analysis.

### Glycolysis measurements with HYlight sensor and analysis

Glycolysis dynamics were measured using a genetically encoded fluorescent biosensor for fructose 1,6-bisphosphate (FBP) termed HYlight as described ^23^. Neurons were transfected with HYlight and imaged at 11-12 DIV. Before the beginning of image acquisition, the neurons were pre-equilibrated with Tyrode buffer perfusion for 10 minutes in the stimulation chamber. In 2-deoxyglucose (2DG) experiments, 5mM glucose in the Tyrode buffer was replaced with 5mM 2DG as indicated with the arrow in Figure 4D. For each imaging session, first pre-stimulus fluorescence baseline (R_0_) measurements were performed for 5 minutes in the absence of electrical activity as a control, then action potentials were evoked by field stimulation (100mA,10 Hz,600 AP). Time-lapse movies from neurons expressing HYlight were acquired as described previously ^23^ for 15-20min. Briefly, the neuron images were acquired for each time point by using 488 nm and 405 nm lasers for excitation and emission detection ranges at 492-598 nm and 490-596 nm, respectively. HYlight fluorescence ratio was calculated from individual neurons by selecting at least 3 ROIs with an average area of 20-30 µm^2^ from both proximal and medial/distal (>75 µm away from soma boundary) neuronal processes. Then the normalized HYlight emission induced by 488nm and 405nm excitations (ΔR/R_0_) was calculated as a readout of temporal changes in FBP level by dividing the HYlight intensity data points with pre-stimulus intensity.

### ATP level imaging and analysis

Single-wavelength genetically encoded ratiometric fluorescent sensor mRuby-iATPSnFR^1.0^ was used for cytoplasmic ATP measurements (Synapsin-cyto-mRuby3-iATPSnFR^1.0^) ^62^. Neurons were transfected with mRuby-iATPSnFR^1.0^ and Mito-BFP plasmids. Time-lapse movies were acquired from 11-12 DIV neurons for a total of 10-15 minutes at 0.1-0.2 Hz using 488 nm, 568nm, and 405 nm lasers for excitation. O-GlcNAc transferase activity was blocked by 5 µM OSMI-4 treatment. Neurons were preincubated with OSMI-4 for 30 minutes before imaging, and perfused with Tyrode buffer containing OSMI-4 during live-cell imaging. In 2-deoxy-D-glucose (2DG) and Oligomycin experiments, 5mM glucose in the Tyrode buffer was replaced with 5mM 2DG and 2 µM Oligomycin added during live-cell imaging. For each imaging session, first pre-stimulus fluorescence baseline (F_0_) measurements were performed for 2 minutes in the absence of electrical activity as a control, then action potentials were evoked by field stimulation (100mA,10 Hz,600 AP). Changes in mRuby-iATPSnFR^1.0^ fluorescence (ΔF/F_0_) were calculated by dividing the fluorescent intensity differences between frames with pre-stimulus intensity for each stimulation. For each neuron, one to four regions of interest (ROIs) with an average area of 20-30 µm^2^ of neuronal processes (<75 µm away from the soma) were selected for image analysis. F/F_0_ for mRuby and iATPSnFR^1.0^ channels were plotted independently without the ratiometric calculations for each condition for the mRuby-iATPSnFR^1.0^ sensor data.

### Mitochondrial membrane potential measurements

Primary hippocampal neuron cultures (9-13 DIV) were treated with 2 µM Thiamet-G or vehicle DMSO (Sigma-Aldrich) overnight. For mitochondrial membrane potential measurements, neurons were stained with tetramethylrhodamine methyl ester (TMRM) (Invitrogen) at a non-quenching concentration (20nM), and co-stained with membrane potential insensitive mitochondria dye MitoTracker Green FM (Invitrogen) (40nM) for 20 minutes. For live-cell imaging, coverslips were transferred to Hibernate E (BrainBits) medium containing 5nM TMRM as described previously ^60^. Live-cell imaging was performed at 37 ºC, using Zeiss LSM 780 laser scanning confocal microscope equipped with a temperature-controlled stage, and Plan-Apochromat 100x/1.40 Oil DIC M27 objective. For the relative quantification of membrane potential for each mitochondrion, the fluorescence intensity of TMRM was normalized to the MitoTracker Green signal for each mitochondrion from neuronal processes.

### Immunocytochemistry of neurons

11-15 DIV disassociated primary neuron cultures were fixed and immunostained immediately after indicated neuronal stimulations. Neuronal activity was manipulated either with field stimulation (100mA,10 Hz,600 AP) or 6 hours of 50 µM picrotoxin (PTX, Tocris) treatment ^65^, either alone or with 10 µM 2,3-Dioxo-6-nitro-1,2,3,4-tetrahydrobenzo[f]quinoxaline-7-sulfonamide (NBQX, Tocris). Neurons were fixed with 4% PFA and 4% sucrose in PBS for 10 minutes at room temperature, and immunostained with the antibodies in 1x GBD (10% goat serum, 1% bovine serum albumin, and 0.1% Triton X-100 in PBS) as previously described ^66^. The coverslips were mounted using Fluoromount-G (Southern Biotech, Birmingham, AL). The following primary antibodies were used for staining: mouse anti-O-GlcNAc (RL2, Abcam), chicken Anti-βIII Tubulin (Tuj1) (Novus Biological), rabbit anti-Tomm20 (Sigma-Aldrich). The secondary antibodies were: goat anti-mouse Alexa 488 (Life Technologies A11029), goat anti-rabbit Alexa 568 (Life Technologies A11036), goat anti-chicken Alexa 647 (Life Technologies A21449), goat anti-chicken Alexa 405 (Abcam), and donkey anti-mouse Alexa 647 (Jackson ImmunoResearch Laboratories, Inc.). Images were acquired using a Zeiss LSM780 confocal laser scanning microscope, Plan-Apochromat 100x/1.40 Oil DIC M27 objective using the same image acquisition settings across different conditions and coverslips. DAPI staining was used to define the perinuclear regions in the soma compartment for each neuron (Figure 2). Axonal and dendritic neuronal processes are determined based on their mitochondrial and structural morphology as previously described ^67,68^. In Figure 3, the mitochondrial area was defined based on the Anti-Tomm20 positive areas with a mean gray value of 25 and higher. The images were further analyzed using Fiji ^64^, using only linear adjustments of brightness and contrast for visualization.

### Preparation of mitochondrial fractions and protein analysis

Mitochondrial fractions were prepared from 11-15 DIV primary cortical neuron cultures, plated on 6-well plates, by homogenization in mitochondrial isolation buffer (MIB) (10mM Tris-HCl (pH 7.4), 10mM KCl, 250mM Sucrose, 1 mM EDTA, 1x Protease Inhibitor Cocktail Set III (Calbiochem), 0.1mM PMSF, 2 µM Thiamet-G and 2mM DTT) and differential centrifugation. Each well containing neurons was washed (∼1.5 × 10^7^ cells per condition) and incubated with freshly prepared MIB on ice for 10 minutes. Cells were detached with a cell scraper and homogenized (20-30 strokes with a tight-fitting B pestle in a 1ml Dounce homogenizer). Nuclear waste and large debris were pelleted by centrifugation at 700g for 10min. The supernatant (containing mitochondria) was centrifuged again at 10,000g for 10min to pellet the crude mitochondrial fraction. The supernatant (cytoplasmic fraction) was collected and concentrated using Amicon Ultracel-10 centrifugal filter units (Millipore). 20% of mitochondrial and 10% of cytoplasmic fractions were then loaded in SDS-PAGE and analyzed by western blotting. The whole-cell lysate was collected using lysis buffer (50mM Tris-HCl (pH 7.4), 150 mM NaCl, 1 mM EDTA (pH 8.0), and 2% NP-40 with 1x Protease Inhibitor Cocktail Set III, 0.1 mM PMSF, 2 µM Thiamet-G and 2 mM DTT), centrifuged at 13,500g for 15min to remove cell debris and analyzed by western blotting. The following antibodies were used for probing blots: mouse anti-O-GlcNAc (RL2, Abcam), rabbit anti-ATP5β (Sigma-Aldrich), rabbit anti-GPI (Thermo-Fisher), mouse anti-actin (Sigma-Aldrich), rabbit anti-OGT (DM-17, Sigma-Aldrich), rabbit anti-OGA (MGEA5) (Proteintech), mouse anti-tubulin (Sigma-Aldrich), rabbit anti-RHOT1(Miro1) (Aviva Systems Biology), mouse anti-cytochrome c (Abcam), rabbit anti-TFAM (mtTFA) (Abcam), mouse anti-Slc25a4/5 (ANT1/2) (Abcam), rabbit anti-Suclg1 (Cell signaling Technology), goat anti-mouse peroxidase (Jackson Immuno Research Laboratories, Inc.), goat anti-rabbit peroxidase (Jackson Immuno Research Laboratories, Inc.). For quantitative western blot measurements, image exposure times were optimized for the linear range of detection using the Azure Biosystem gel documentation system. The images were further analyzed using the Fiji gel analyzer ^64^, applying only linear adjustments of brightness and contrast for visualization.

### Respirometry measurements with primary cortical neurons

Oxygen consumption and extracellular acidification rates were measured using an Agilent Seahorse XFe96 Analyzer and Seahorse XF Mito Stress Test protocol, from 12-15 DIV primary cortical neurons. Neurons were cultured on XF96e plates at 45,000/well density and treated with 1 µM Ara-C (Cytarabine, Tocris) for two days at 3 DIV as described above. Neuronal activity was manipulated with 6 hours of 50 µM picrotoxin (PTX, Tocris) treatment, either alone or with 5 µM OSMI-4 as demonstrated in Figure S3. The neurons were treated with OGA inhibitor 5 µM Thiamet-G or DMSO (Control) and incubated overnight for the experiments to upregulate O-GlcNAcylation. For respirometry measurements, the neuronal maintenance medium was exchanged with XF DMEM Base Medium (pH 7.4) with no phenol red (Agilent) supplemented with 5 mM glucose and 1 mM pyruvate. NucBlue dye (Thermo Fisher Scientific) was included to stain nuclei for cell counting immediately after the assay. Respiration was measured under basal conditions as well as after injections of 1 µM Oligomycin (Sigma-Aldrich), 0.5 µM FCCP (Sigma-Aldrich), and 0.5 µM Rotenone (Sigma-Aldrich)/0.5 µM Antimycin A (Sigma-Aldrich) injections. Basal and maximal respiration, reserve capacity, basal glycolysis and glycolytic capacity values were calculated as previously described ^69^ after cell count normalization with FluxNorm (see details at https://github.com/pekkurnazlab/FluxNormalyzer). After each assay, the neuronal enrichment percentage was calculated by immunostaining of seahorse plates with neuron-specific antibody (anti-NeuN) and cellular nuclei (NucBlue).

### Mass spectrometry sample preparation and analysis

#### O-GlcNAcylated protein enrichment and in-gel digestion

Mitochondrial fractions were prepared from 11-15 DIV primary cortical neuron cultures, treated either with 5 µM Thiamet-G or vehicle overnight. Mitochondrial pellets were dissolved in the lysis buffer. 9 µg of succinylated Wheat Germ Agglutinin (sWGA)-bound agarose beads (Vector Laboratories, Burlingame, CA) and mitochondrial lysates were incubated for 12hr at 4 ºC for O-GlcNAcylated protein enrichment. Monosaccharide inhibitor GlcNAc (40mM) was added before the incubation of mitochondrial lysate with sWGA beads as a negative control. To eliminate non-specific binding sWGA beads were washed with lysis buffer 4 times with gentle agitation and resuspended with 1x Laemmli buffer. 80-90% of each sample was separated by pre-cast 4-20% SDS-PAGE (Bio-Rad). “SimplyBlue SafeStain” (Invitrogen) stained gel bands were excised, minced, and prepared for mass spectrometry analysis as previously described ^70^. Briefly, the gel slices were cut into 1mm x 1 mm cubes and de-stained 3 times by first washing with 100 µl of 100 mM ammonium bicarbonate for 15 minutes, followed by additions of the same volume of acetonitrile (ACN) for 15 minutes. The samples were dried in a speedvac and reduced by mixing with 200 µl of 100 mM ammonium bicarbonate-10 mM DTT and incubated at 56 °C for 30 minutes. The solution was removed, and 200 μL of 100 mM ammonium bicarbonate-55mM iodoacetamide was added to gel pieces and incubated at room temperature in the dark for 20 minutes. After removing the supernatant and washing the gel pieces with 100 mM ammonium bicarbonate for 15 minutes, the same volume of ACN was added to dehydrate the gel pieces. The samples were dried in a speedvac and processed for trypsin digestion. For digestion, ice-cold trypsin (0.01 µg/µL) in 50 mM ammonium bicarbonate was added to cover the gel pieces and incubated on ice for 30 min. After complete rehydration, the excess trypsin solution was removed, replaced with fresh 50 mM ammonium bicarbonate, and left overnight at 37°C. The peptides were extracted twice by the addition of 50 µl of 0.2% formic acid and 5 % ACN and vortex mixing at room temperature for 30 min. The supernatant was removed and saved. A total of 50 µl of 50% ACN-0.2% formic acid was added to the sample, which was vortexed again at room temperature for 30 min. The supernatant was removed and combined with the supernatant from the first extraction for further analysis.

#### Identification of O-GlcNAc captured proteins

The samples were analyzed by ultra-high pressure liquid chromatography (UPLC) coupled with tandem mass spectroscopy (LC-MS/MS) using nano-spray ionization. The nanospray ionization experiments were performed using an Orbitrap fusion Lumos hybrid mass spectrometer (Thermo) interfaced with nano-scale reversed-phase UPLC (Thermo Dionex UltiMate™ 3000 RSLC nano System) using a 25 cm, 75-micron ID glass capillary packed with 1.7-µm C18 (130) BEHTM beads (Waters Corporation). Peptides were eluted from the C18 column into the mass spectrometer using a linear gradient (5–80%) of ACN at a flow rate of 375 µl/min for one hour. The buffers used to create the ACN gradient were: Buffer A (98% H2O, 2% ACN, 0.1% formic acid) and Buffer B (100% ACN, 0.1% formic acid). Mass spectrometer parameters are as follows; an MS1 survey scan using the orbitrap detector set to (mass range (m/z): 400-1500 (using quadrupole isolation), 120000 resolution setting, spray voltage of 2200 V, Ion transfer tube temperature of 275 C, AGC target of 400000, and maximum injection time of 50 ms) was followed by data-dependent scans (top speed for most intense ions, with charge state set to only include +2-5 ions, and 5 second exclusion time, while selecting ions with minimal intensities of 50000 at in which the collision event was carried out in the high energy collision cell (HCD Collision Energy of 30%). The fragment masses were analyzed in the ion trap mass analyzer (With the ion trap scan rate of turbo, first mass m/z was 100, AGC Target 5000, and maximum injection time of 35ms). PEAKS Studio 8.5 software was used for peptide identification and label-free quantification of the relative abundance of all peptides detected in Thiamet-G or vehicle samples ^71^.

#### LC-MS/MS data analysis

Label-free quantification data for the O-GlcNAc captured proteins from paired Thiamet-G or vehicle treated primary neuron cultures were generated in three separate experiments. These values represent an estimated abundance for each protein in each sample. Prior to merging the sample data, proteins with < 2 peptides and/or not detected in at least one sample in each of the groups (Thiamet-G and vehicle treated) were removed. The merged data matrix was used as input to the Perseus software package ^72^, which performed a series of processing steps. First, label-free quantification values were log transformed. Second, abundance values were imputed for proteins in samples where the protein was undetected. The imputed values were randomly assigned from a gaussian distribution with the Perseus default parameters for width and downshift. Finally, the processed abundance value matrix was imported into the R statistical computing environment (www.cran.org) for additional analyses. To identify differentially abundant proteins, moderated t-tests from the limma package ^73^ were used to compare the two groups (Thiamet-G treatment vs vehicle treatment) where the experimental design was modeled upon treatment (∼0 + treatment) and p-value < 0.05 was used as the cutoff for significance. Scatter plots and pie charts were made with the ggplot2 package for R ^74^. Of primary importance in this study were proteins detected in the mitochondrial databases MitoMiner, Mouse Mitocarta 3.0 ^25,26^ and the O-GlcNAc Database ^27,75^. Proteins identified by mass spec were annotated with the information from these databases including subcellular localization, mitochondria-specific localization, and mitochondria-related processes. Annotation data for the identified proteins were used for filtering to relevant subsets, aggregated, and used as input to generate pie charts and other visualizations. Enrichment analysis was performed with gprofiler2 ^76,77^ within R using a p-value cutoff of 0.05. Network analysis was performed using the string-db.org web application ^78^ and exported to Cytoscape ^79^ for image preparation.

### Statistical tests

Throughout the paper, error bars indicate the mean ± SEM unless otherwise noted. All p-values and the number of replicas were indicated in the figure legend for each experiment. Statistical analysis was performed with GraphPad Prism 8 for MacOSX or R. The Mann-Whitney U test was used to determine the significance of differences between the two conditions. Multiple conditions were compared by the Kruskal-Wallis nonparametric ANOVA test, which was followed by Dunn’s multiple comparisons test or by one-way ANOVA with post hoc Tukey’s test as appropriate to determine the significance of differences across every condition to control condition.

## Notes

### Competing Interest Statement

The authors have declared no competing interest.

